# Integrating Subclonal Response Heterogeneity to Define Cancer Organoid Therapeutic Sensitivity

**DOI:** 10.1101/2021.10.15.464556

**Authors:** Jeremy D. Kratz, Shujah Rehman, Katherine A. Johnson, Amani A. Gillette, Aishwarya Sunil, Peter F. Favreau, Cheri A. Pasch, Devon Miller, Lucas C. Zarling, Austin H. Yeung, Linda Clipson, Samantha J. Anderson, Alyssa K. DeZeeuw, Carley M. Sprackling, Kayla K. Lemmon, Daniel E. Abbott, Mark E. Burkard, Michael F. Bassetti, Jens C. Eickhoff, Eugene F. Foley, Charles P. Heise, Randall J. Kimple, Elise H. Lawson, Noelle K. LoConte, Sam J. Lubner, Daniel L. Mulkerin, Kristina A. Matkowskyj, Cristina B. Sanger, Nataliya V. Uboha, Sean J. Mcilwain, Irene M. Ong, Evie H. Carchman, Melissa C. Skala, Dustin A. Deming

## Abstract

Tumor heterogeneity is predicted to confer inferior clinical outcomes, however modeling heterogeneity in a manner that still represents the tumor of origin remains a formidable challenge. Sequencing technologies are limited in their ability to identify rare subclonal populations and predict response to the multitude of available treatments for patients. Patient-derived organotypic cultures have significantly improved the modeling of cancer biology by faithfully representing the molecular features of primary malignant tissues. Patient-derived cancer organoid (PCO) cultures contain numerous individual organoids with the potential to recapitulate heterogeneity, though PCOs are most commonly studied in bulk ignoring any diversity in the molecular profile or treatment response. Here we demonstrate the advantage of evaluating individual PCOs in conjunction with cellular level optical metabolic imaging to characterize the largely ignored heterogeneity within these cultures to predict clinical therapeutic response, identify subclonal populations, and determine patient specific mechanisms of resistance.

## Introduction

Cancer cell heterogeneity based on the molecular profile and metabolic state is commonly identified in research settings. However, limited clinical means exist to identify this heterogeneity and it is largely unknown how to use this information in predicting treatment response or guiding future therapeutic strategies. Patient-derived cancer organoids (PCOs) have become the preferred *in vitro* culture technique for modeling tumor biology across many cancers^1-5^. PCOs are generated with high efficiency, retain the molecular characteristics of their parent tissues, and therapeutic response that correlate with clinical response for patients^6-10^. To date, studies have examined the PCO response on the culture-well level, yet the heterogeneity between individual PCOs is not taken into consideration. Each well can contain hundreds of individual PCOs, each representing distinct clonal or oligoclonal units of the original tumor. We have developed the use of single-cell optical metabolic imaging (OMI) technologies using two-photon microscopy to provide a detailed readout of PCO treatment response within intact, live samples^11-18^. OMI non-invasively measures treatment response without extrinsic reagents (e.g., labels or dyes) and has previously monitored metabolic heterogeneity in multiple cancer models including response predictions in prospective clinical studies^8,19-22^.

Tools capable of capturing heterogeneity are of vital importance when subclonal heterogeneity can confer primary or acquired clinical resistance^23,24^. Epidermal growth factor receptor (EGFR) inhibition, with cetuximab or panitumumab, is a standard treatment for patients with *KRAS, NRAS*, and *BRAF*^*V600*^ wild-type metastatic colorectal cancer (CRC)^25-27^. Many mechanisms of resistance to EGFR inhibition have been described including metabolic changes, clonal evolution, and the development of subclonal heterogeneity^24,28-30^. Acquired mechanisms of resistance following EGFR inhibition cannot be predicted for individual patients from pretreatment bulk sequencing or circulating tumor DNA (ctDNA) alone. The selective pressure of EGFR inhibition determines clones that either develop activating alterations in RAS signaling^31-34^ or differential expression of key mediators across multiple pathways^35-39^. Clinical ctDNA can detect the acquisition of alterations associated with resistance to EGFR inhibition, but these most often are found at extremely low allele frequencies and cannot track important metabolic changes that occur in response to therapy^40^. The metabolic and molecular heterogeneity that develops in response to EGFR inhibition in CRC makes this an ideal clinical scenario to evaluate the use of PCOs to characterize this heterogeneity.

Here, we assess individual PCO response to understand the contributions of subclonal populations in assaying the responsiveness to chemotherapy and radiation across a diverse set of advanced cancers. Treatment response and organoid viability are determined using single-cell OMI and change in individual organoid diameter^15^. These imaging analyses determine the response to treatment across the population of PCOs, rather than current techniques of culture well-level assessments, to recognize subclonal populations that predict clinical response to therapy. Furthermore, CRC PCOs are treated with escalating doses of panitumumab over time until the PCOs develop resistance, which provides a potential marker of treatment response durability and a framework to identify patient specific resistance mechanisms using organoid DNA and RNA sequencing. Overall, we demonstrate that PCOs can predict clinical activity of therapeutics across diverse treatments, identify clinically relevant heterogeneity using morphologic and metabolic imaging, and identify patient-specific resistance mechanisms.

## Methods

### Cell isolation and organotypic culture techniques

All studies were completed following Institutional Review Board (IRB) approval with informed consent obtained from subjects through the University of Wisconsin (UW) Molecular Tumor Board Registry (UW IRB#2014-1370) or UW Translational Science BioCore (UW IRB#2016-0934). Briefly, tissue obtained from needle, endoscopic biopsy, or primary surgical resection was placed in chelation buffer for at least 30 minutes. Two PBS washings were performed for tissues acquired using endoscopic biopsies. Digestion was performed in DMEM stock (**Table S1**) with the addition of collagenase (50 mg/mL) and dispase (10 mg/mL). Tissue was digested with intermittent shaking and mechanical disruption using a p1000 pipette from 15 minutes to 2 hours dependent upon tissue size. Malignant fluids (ascites, pleural effusion, or pericardial effusions) were initially pelleted and separated using a Ficoll solution preparation (Sigma). Once digestion or Ficoll preparation was completed, the samples were centrifuged at 1000 rpm at 4°C for 5 minutes and resuspended in ADF stock (**Table S1**). PCO suspensions were immediately mixed at a 1:1 ratio with Matrigel matrix (Corning). Droplet suspensions were plated and set for three to five minutes at 37°C then inverted for at least 30 minutes to solidify the matrix and avoid the direct interface of PCOs with the plastic interface. Plated cultures were overlaid with 450μL of feeding medium supplemented with tissue dependent components (**Table S1**) and incubated at 37°C in 5% CO_2_. Medium was replaced every 48-72 hours.

### Therapeutics studies

PCOs were collected from 24-well culture plates and passaged to single well 35mm glass dishes (Cellvis) or 24-well glass plates. Images were taken on a Nikon Ti-S inverted microscope using a 4x objective prior to treatment. Pharmacologic agents were prepared at physiologic C_max_ including 5-fluorouracil (5-FU) [10 μM]^41^, oxaliplatin [5 μM]^42^, SN-38 [1.5 nM]^43^, gemcitabine [50 μM]^44^, paclitaxel [50 nM]^44^, olaparib [200 nM]^45^, osimertinib [30 nM]^46^, and panitumumab [1.6 μM]^47^ (**Table S2**). The duration of therapy was continuous over 48 hours for all agents with the exception of gemcitabine which was dosed over 24 hours to model clinical pharmacokinetics^48^. Given the uncertainty of monoclonal antibody (mAb) delivery at the time of study design, EGFRi assays were performed after 1-hour of preincubation of the PCOs that were suspended in physiologic panitumumab (**Fig S1**). Pharmacologic agents were received from the University of Wisconsin Carbone Cancer Center Pharmacy including 5-FU (Fresenius), gemcitabine (Hospira), oxaliplatin (Hospira), paclitaxel (Athenex), and panitumumab (Amgen). Additional agents prepared from stock powders including SN-38 (Sigma), olaparib (LC Laboratories), and osimertinib (LC Laboratories). DMSO concentration was <0.1% v/v for all final culture conditions. Radiation was delivered using XStrahl RS-225 cabinet delivering a dose rate of approximately 3.27 Gy/minute using an aluminum filter with monthly quality assurance performed by the UW Medical Radiation Research Calibration Lab. After radiation, media was exchanged at 48 hours and response relative to control was assayed at 96 hours. All therapeutic studies were performed as three independent biological replicates.

### EGFR inhibition with dose escalation studies

Following maturation, PCOs were treated with stepwise dose escalation. Cultures were started at 20% physiologic C_max_ panitumumab (46ug/ml) and assessed for growth (normalized change in diameter at 96 hours). If cultures surpassed ≥20% growth, stepwise dose escalation was repeated with an interval increase of 20% C_max_. Time to resistance (TTR) was defined by duration of time from the initiation of dose escalation to persistent growth at 100% C_max_ panitumumab. Epifluorescence signal was collected using a Nikon Ti-S epifluorescence microscope equipped with a 4X air objective (Nikon CFI Plan Fluor, NA 0.13, FOV: 5.5mm x 5.5mm) and CMOS camera (Flash4, Hamamatsu). NAD(P)H fluorescence was excited with a DAPI filter cube (Nikon, ex: 361-389nm / em: 435-485nm, integration time 100 ms) and FAD fluorescence was excited by Intensilight C-HGFI (Nikon) through a custom filter set (Semrock, ex: 426-486nm / em: 525.5-630.5nm, integration time 200 ms). Epifluorescence signal was thresholded in ImageJ based on NAD(P)H signal intensity to define organoid-level regions of interest (ROI). ROIs were used to measure the mean intensity for the optical redox ratio, defined as the intensity of NAD(P)H/FAD.

### Brightfield and epifluorescence image analysis

Normalized change in diameter as an assessment of growth was measured from brightfield images collected at 0 hours and 48 hours using ImageJ (NIH) and converted from pixel to distance using the triplicate measurements of a 1000μm standard. The distance to the edge of the Matrigel was calculated from the closest edge of the PCO with a perpendicular axis to the matrix edge to represent the shortest length. Experimental correlations between baseline size, relative passage number and PCOs per field of view were compared to growth using the adjusted coefficient of determination (R^2^) by OriginPro 2020. Effect size was determined using Glass’s Delta (GΔ) which normalizes the difference in mean observed signal to the control population normalized to the standard deviation of the control^49^. Interquartile ranges were compared using effect size normalized to the first quartile. Epifluorescence signal was thresholded in ImageJ to NAD(P)H signal intensity to define organoid-level regions of interest (ROIs). The ROIs were then used to measure the mean intensity for the organoid-level optical redox ratio (ORR) defined as intensity ratio of NAD(P)H/FAD.

### Two-photon optical metabolic imaging (OMI) acquisition and analysis

Briefly, NAD(P)H and FAD were excited at 750 nm and 890 nm, respectively, using a tunable Ti:sapphire laser (Coherent, Inc), an inverted microscope (Nikon, Eclipse Ti), and a 40x water immersion (1.15NA, Nikon) objective. Both NAD(P)H and FAD images were obtained for the same field of view. FAD fluorescence was isolated using an emission bandpass filter of 550/100 nm, while NAD(P)H fluorescence was isolated using an emission bandpass filter of 440/80 nm. Fluorescence lifetime data was collected using time-correlated single-photon counting electronics (SPC-150, Becker and Hickl) and a GaAsP photomultiplier tube (H7422P-40, Hamamatsu). Images (512 × 512 pixels) were obtained using a pixel dwell time of 4.8μs over 60s total integration time. A Fluoresbrite YG microsphere (Polysciences Inc.) was imaged as a daily standard for fluorescence lifetime. The lifetime decay curves were fit to a single exponential decay, and the fluorescence lifetime was measured to be 2.1 ns (*n* = 7), which is consistent with published values ^19,50,51^.

Fluorescence lifetimes were extracted using SPCImage software (SPCImage v8.1, Becker & Hickl). A bin of 3×3 pixels was used to maintain spatial resolution, the fluorescence lifetime decay curve was convolved with the instrument response function and fit to a two-component exponential decay model using,

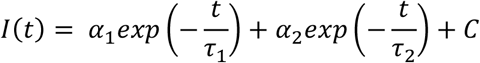

where *I*(*t)* is the fluorescence intensity at time *t, α* is the fractional contribution of each component, *τ* is the lifetime of each component, and C accounts for background light. The two lifetime components are used to distinguish between the free and bound forms of NAD(P)H and FAD^52,53^. The mean fluorescence lifetime was calculated using, *τ*_*m*_ = *α*_1_*τ*_1_ + *α*_2_*τ*_2_. The decay curves for NAD(P)H and FAD were integrated for each pixel to obtain intensity values. The optical redox ratio (ORR) was calculated by dividing the intensity of NAD(P)H by the intensity of FAD. All reported ORRs are normalized to the average control ORR of the same patient and time point.

A semi-automated cell segmentation algorithm was developed using Cell Profiler software^54^. This system identified pixels belonging to nuclear regions using a customized threshold code. Cells were recognized by propagating out from the nuclei within the image. To refine the propagation and to prevent it from continuing into background pixels, an Otsu Global threshold was used. The cell cytoplasm was defined as the cell borders minus the nucleus. Values for NAD(P)H lifetime variables (*τ*_*m*_, *τ*_1_, *τ*_2_, *α*_1_, *α*_2_), FAD lifetime variables, NAD(P)H intensity, and FAD intensity were averaged for all pixels within the cytoplasm of each cell, as described previously^14,55,56^. The fluorescence lifetime redox ratio (FLIRR) was calculated by dividing the fractional contribution of bound NAD(P)H (*α*_*2*_) by the fractional component of bound FAD (*α*_*1*_)^57-60^. The FLIRR enables metabolic comparison of organoids captured on different days as it overcomes several experimental limitations of fluorescence intensity imaging. Fluorescence intensity measurements are influenced by laser power, detector gain, light scattering, and the concentration of the fluorophore; however, fluorescence lifetimes are independent of these experimental factors^57,59^. Although the FLIRR is a useful measurement of cellular metabolism, it is not a substitute for the optical redox ratio and instead represents an additional variable to compare metabolic endpoints^55^. Hierarchical cluster analysis of OMI parameters was completed using group average cluster methods defined using Euclidean distant type (OriginPro 2020).

### Subpopulation modeling in response heterogeneity

The collective cell population of each treatment type was input into a Gaussian mixture distribution model^61^ (MATLAB, version 2018a, MathWorks, Natick, Mass) given by:

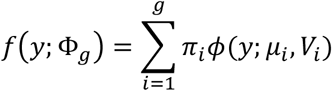

where *g* is the number of subpopulations, *ϕ* (*y*; *μ*_*i*_, *V*_*i*_ ) is the normal probability density function with mean *μ*_*i*_, variance *V*_*i*_, and *π*_*i*_ is the mixing proportion. Goodness of fit was calculated given a set of subpopulations (*g* = 1, 2, or 3) using an Akaike information criterion^62^. The number of subpopulations was determined based on the lowest Akaike score. Probability density functions were normalized to ensure that the area under the curve for each treatment group was equal to 1. Treatment effect size was calculated using GΔ^49^ and defined as meaningful if >0.5 for single cell OMI and >1.5 for sphere growth analysis based on prior work^8^. Due to expected heterogeneity across mixed populations, the pooled analyses of multiple cultures in response to EGFRi was performed by Gaussian population modeling using GraphPad Prism 9.

### Next generation sequencing of PCOs for DNA and RNA

PCOs were collected from ∼4-6 wells, washed in 1x sterile PBS, and stored at -80°C as a cell pellet until DNA and RNA isolation. For RNA collection, following washing, PCOs were stored at -80°C in RNA_later_ (Qiagen) until library preparation. DNA was isolated on a Maxwell 16 AS2000 or Maxwell CSC (Promega) using the Maxwell DNA LEV Blood Kit (Promega #AS1290) according to manufacturer instructions. Libraries were prepared using the QIASeq Human Comprehensive Cancer Panel Kit (Qiagen #333515) and sequenced on an Illumina HiSeq 2500 or NovaSeq 6000. All samples were collected in triplicate biological replicates at baseline and after achieving resistance to EGFRi with persistent growth at 100% physiologic C_max_. RNA libraries were constructed using TruSeq Stranded total RNA with rRNA reduction (Illumina). Quality control was performed by Eukaryote Total RNA electrophoresis using NanoDropOne. The library included 2×150bp paired end reads with sequencing performed by NovaSeq 6000 using MiSeq NanoCell at the UW Biotechnology Center.

### Subclonal analyses

Subclone counts were analyzed by counting the number of unique alterations present at a frequency between 10% and 30%, that were likely to alter the protein (missense mutations, insertions/deletions, and alterations to splice donors/ acceptors), and were not likely to be artifacts of sequencing. Artifacts in this case were defined as alterations that occurred in more than five individual patients. Subclones were analyzed using R version 3.6.1 (R Core Team 2019; https://www.R-project.org) and the tidyr v1.1.1^63^ and dplyr v0.8.5^64^ packages (https://CRAN.R-project.org). Relative variant allele frequency (rVAF) was defined by normalizing to the molecularly informed tumor content provided by the commercial vendor. Subclonal populations were reported in PCOs assuming 100% tumor content in the reads and defined with rVAF of 10-30%.

### Variant calling and mutation analyses

Sequencing analysis through variant calling was performed at the UW Biotechnology Center. Sequence reads were adapted and quality trimmed using Skewer^65^, aligned to Homo sapiens build 1k_v37 using BWA-MEM^25^, and deduplicated using Picard (http://picard.sourceforge.net) and Je^66^. Base quality scores were recalibrated using GATK^67^ and mutations called using Strelka v-2.8.4^68^ without matched controls and annotated using SNPEff^69^. Resulting VCF files were uploaded to the public Galaxy web platform^70^ (usegalaxy.org) and cross-referenced to ClinVar’s publicly available VCF (accessed 2/13/2021) for annotation of predicted clinical response. Mutations were deemed pathogenic based on the ClinVar database labeled as either “Pathogenic” or “Likely Pathogenic.” Additionally, alterations in *APC, KRAS, NRAS, BRAF, PIK3CA, EGFR, SMAD4, TP53, MLH1, MSH2, MSH6, PMS2, CTNNB1*, and *PTEN* were evaluated for potentially pathogenic mutations not yet curated in ClinVar.

### Statistical analysis of RNASeq data

All statistical calculations were performed using custom *Rmarkdown* scripts. The RSEM estimated counts were first filtered by removing Ensembl gene IDs that generated estimate counts of 0 across all samples then mapped to corresponding gene symbols. The pairwise (Resistance vs. Control for each cell line) differential gene expression was calculated using Bioconductor’s *DESeq2* package^71^. For each sample, pathway expression values were calculated for the curated gene sets (c2) from MSigDB (https://www.gsea-msigdb.org/gsea/msigdb/) using *GSVA* and the Poisson model with the *RSEM* estimated counts^72^. Pair-wise differential pathway expression was calculated using *limma*^73^. Using the *aheatmap* function from the *NMF* package^74^, heatmaps were made of the top 20 genes and top 20 pathways, scaling the regularized log2 (genes) or calculated enrichments across rows. Venn diagrams of the significant genes and pathways (p_adj_ < 0.05) were made using *VennDiagram*^75^.

## Results

### Individual PCOs are heterogeneous in growth, metabolism, and therapeutic response

PCOs were developed from a dedicated tissue-procurement program from multiple cancer types and sample sources, including biopsy, fluid, and surgical specimens from patients (**Table S3**). These samples were named using a code: locally advanced (L), metastatic (M), rectal (R), and colon (C). Other samples from the University of Wisconsin’s Precision Medicine Molecular Tumor Board have names starting “MTB” (**Table S3**). PCOs across patients and cancer types are heterogeneous in both morphology and change in diameter when normalized to baseline diameter (**Fig 1a**). PCO metabolism was assessed using the optical redox ratio (ORR, intensity of NAD(P)H/FAD) which revealed intra-organoid heterogeneity with distinct regions of the PCOs harboring differential metabolic activity even prior to any treatments (**Fig 1b**). Intra-organoid heterogeneity in the intrinsic auto-fluorescence of NAD(P)H and FAD lifetime and ORR was observed across representative PCOs and persisted across multiple cultures (**Fig S2**).

**Figure 1:**
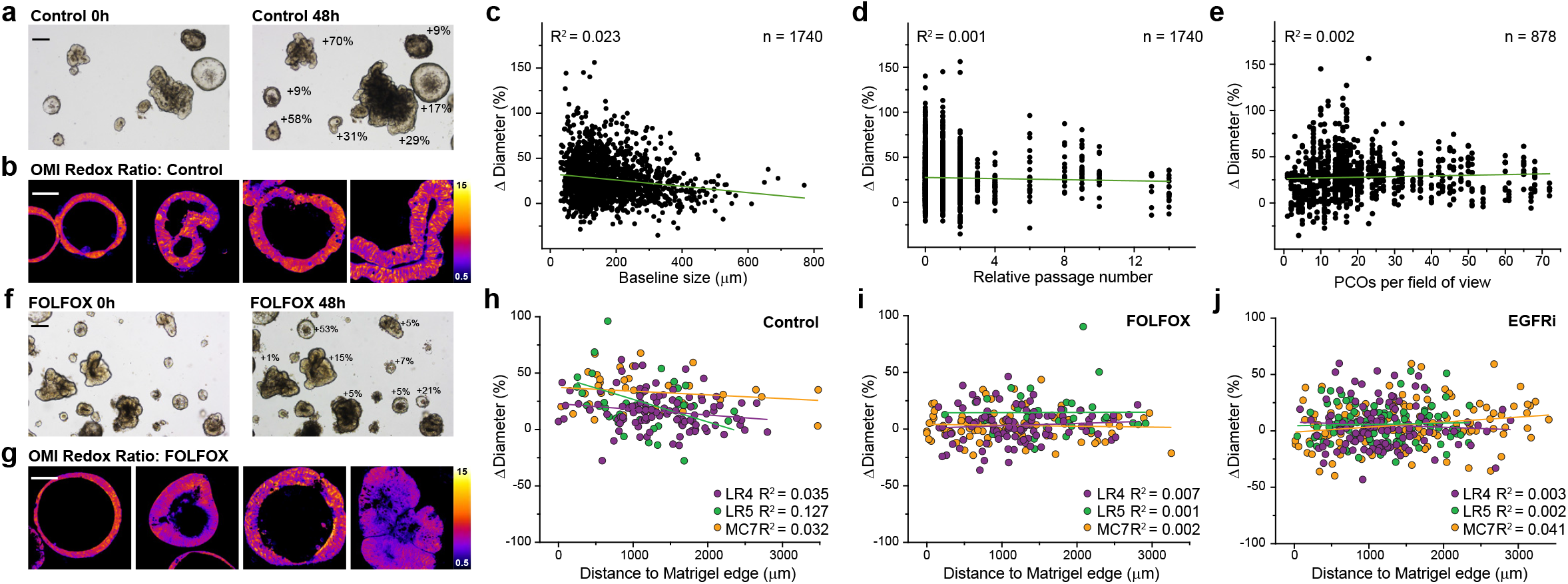
PCO growth and metabolic heterogeneity. (**a**) Representative PCO brightfield microscopy (patient MC1, see **Table S2**) annotated with percent difference in individual organoid diameter and (**b**) representative images of two-photon optical redox ratio (ORR) at 48h for control. (**c**) Correlation plots of baseline size, (**d**) relative passage number, and PCOs per field of view (**e**) against change in diameter (Δ diameter) along with the coefficient of determinant (R^2^). (**f**) Representative therapeutic study (patient MC1) with FOLFOX imaged with brightfield microscopy at 0h and 48h annotated with percent difference in individual organoid diameter and (**g**) representative images of two-photon ORR at 48h. Comparison of PCO location perpendicular to the matrix edge correlated against growth of (**h**) control, (**i**) FOLFOX and (**j**) EGFRi panitumumab across three independent cultures (patients LR4, LR5, MC7, see **Table S2**) along with the coefficient of determinant (R^2^). Scale bars for brightfield (black bar) represent 200μm, scale bars for OMI (white bar) represent 100μm.

### PCO heterogeneity is not driven by organoid size, passage, plating density, or location within the matrix

To determine if culture conditions could be responsible for the heterogeneity observed, multiple baseline experimental parameters were examined. A pooled analysis was performed from 18 patients with CRC that included 22 unique cultures and 1740 individual PCOs. The normalized change in diameter was not predicted by baseline diameter (R^2^= 0.023), relative passage number (R^2^=0.001), or the organoid density measured as the number of PCOs per field of view (R^2^= 0.002) (**Fig 1c-e**). The contributions of baseline diameter (GΔ= 0.32), relative passage number (GΔ= 0.19), and PCOs per field of view (GΔ= -0.32) were minimal across the population (**Fig S3a-c**). Additionally, plating density did not contribute to significant differences in the PCO growth, as only modest changes in median growth were observed compared to the variance within each culture (**Fig S3d-e**). With the combination of 5-FU and oxaliplatin treatment (FOLFOX), variation in individual organoid growth and single cell metabolism persisted within and across treatment groups (**Fig 1f-g**). The location of individual organoids within the 3D matrix did not predict differential growth, therapeutic response to FOLFOX or single agent EGFR inhibitor (**Fig 1h-j**). Additionally, there were no trends by location of organoids with respect to the edge of the matrix in conferring differential growth (GΔ=0.39) or therapeutic response to FOLFOX (GΔ=-0.19) or EGFR inhibitor (GΔ=-0.27) (**Fig S3f-h**).

### PCOs recapitulate driver and subclonal molecular heterogeneity

To evaluate the mutation profile and potential for subclonal populations 34 unique PCO cultures were generated across cancers from 31 subjects. Across the PCO lines, 45 driver alterations were detected. There was 93% concordance for pairwise sets comparing next-generation sequencing (NGS) from clinical specimens and expanded PCOs (**Fig 2a**). The small discordance here is similar to the discordance seen when sampling a single patient’s cancer in separate regions^76,77^. Subclonal mutations were commonly identified within PCOs (**Fig 2b-f**). In PCOs collected from two sites within the same tumor, subclonal alterations largely overlapped, though additional subclonal alterations were seen in the PCOs from one sample (**Fig 2f**).

**Figure 2:**
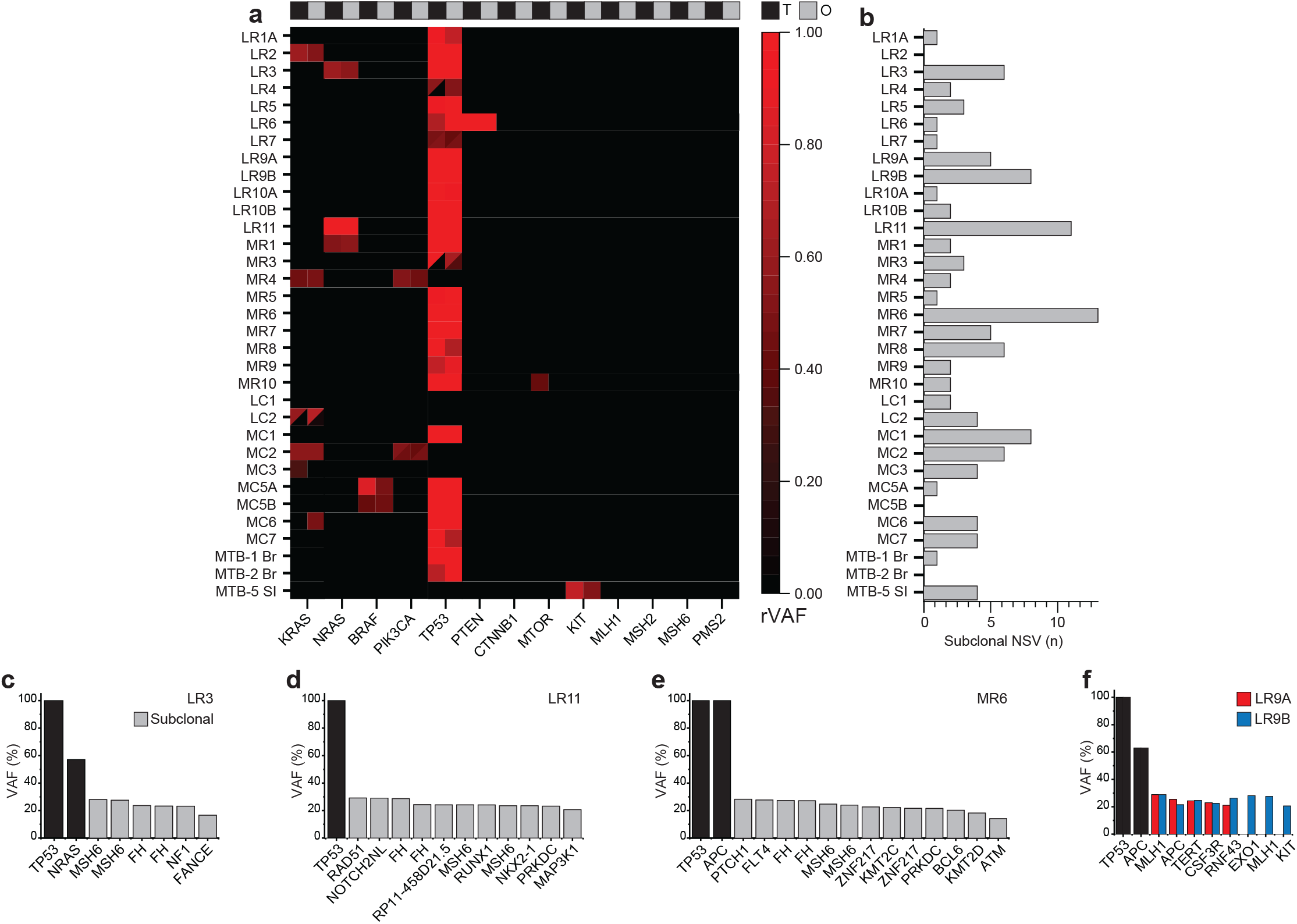
Subclonal molecular heterogeneity in PCOs. (**a**) Heatmap of pathologic alterations between pairwise tumor (black, T) and expanded PCOs (gray, O) for reported pathologic driver alterations with split designation (triangles) for multiple concurrent gene alterations. All variants colored on heatmap according to relative variant allele frequency (rVAF). (**b**) Analysis of subclonal non-synonymous variants (NSV) defined between 10-30% from cancer hotspot testing in PCOs. (**c-e**) Representative examples of clonality including clonal (black) and subclonal variance (gray) across representative cancers. (**f**) Multisite sampling from patient LR9 from endoscopic biopsy with VAF plotted including clonal (black) and discordant subclonal alterations in *EXO1, MLH1*, and *KIT* from LR9A (red) and LR9B (blue).

### PCO growth and metabolic imaging independently track organoid heterogeneity

A total of 2096 individual control organoids were analyzed for change in diameter (**Table S3**). These organoids were derived from cancers of patients with metastatic disease (62%, n = 24), locally advanced rectal cancers (36%, n = 14), and select cases of advanced malignancy at the time of sampling for comprehensive molecular profiling by NGS (**Fig 3a**). The range of mean growth rates observed for cultures (2–54%) was exceeded by the wide range of growth within a given culture (20–201%). All cultures were confirmed to have growth for inclusion of analysis, yet individual cultures harbored a diversity of growth between individual PCOs (**Fig 3b**). PCOs were imaged with 2-photon OMI to evaluate the metabolic heterogeneity of single cells within organoids from 31 primary cultures across 29 independent subjects. Interestingly, there was not a correlation between FLIRR (NAD(P)H α_2_/ FAD α_1_) and the organoid growth rate (R^2^=-0.034), indicating that metabolic imaging provides distinct and potentially complementary information to the change in size measurements (**Fig S4**).

**Figure 3:**
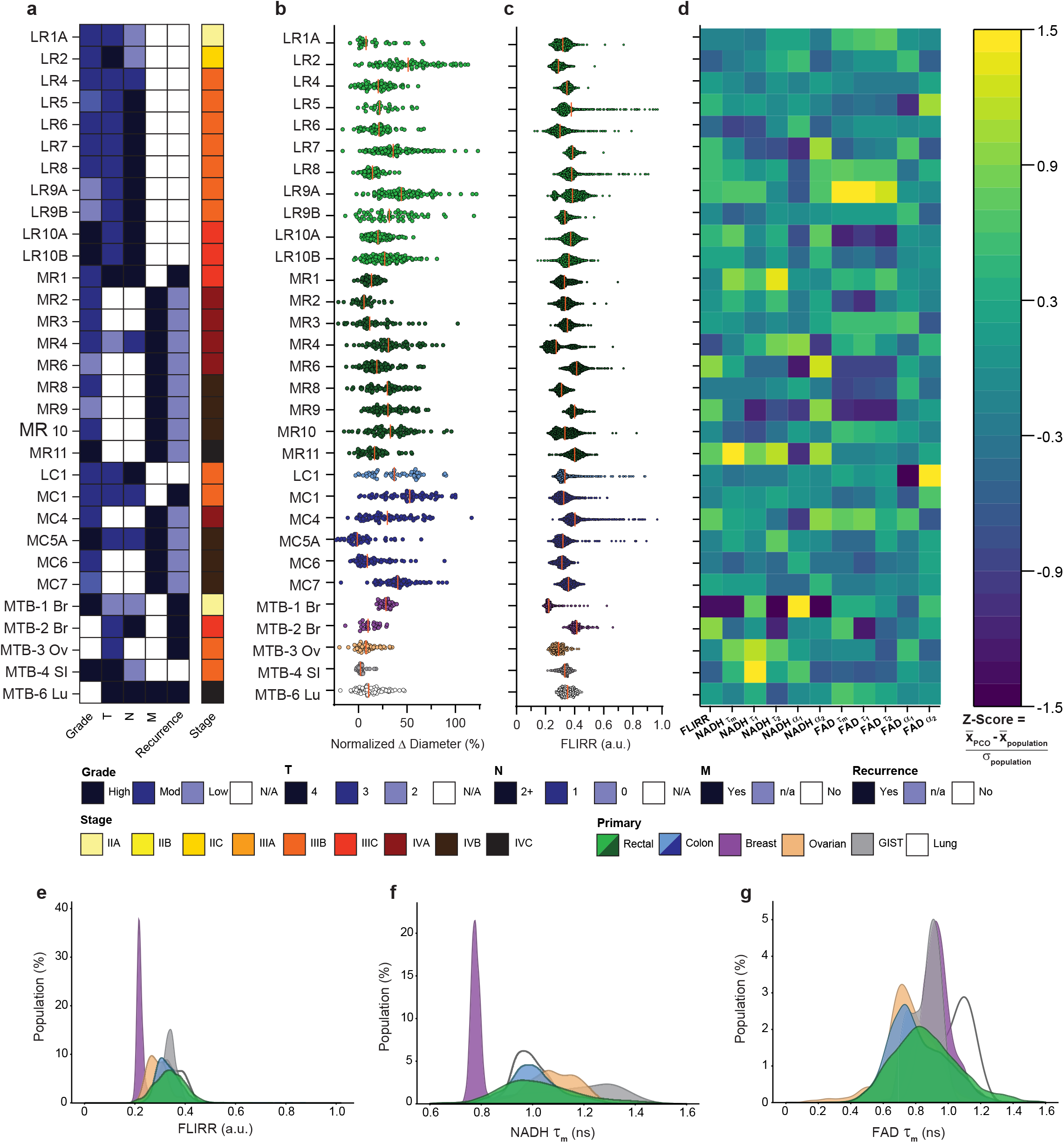
PCO heterogeneity in growth and metabolism as tracked by OMI. (**a**) Baseline clinical characteristics for cultures colored by the key below, including pathologic review of tumor grade and clinical parameters of stage from tumor (T), node (N) and metastases (M) combined to complete diagnostic staging (Stage). Current disease status noted (Recurrence in black). (**b**) Dot plots colored by the key below for primary tumor type (Primary) for localized rectal (light green), metastatic rectal (dark green), localized colon (light blue), metastatic colon (dark blue), breast (purple), ovarian (beige), gastrointestinal stromal tumor (GIST, gray), and lung (white) with each point representing an individual organoid for normalized Δ diameter at 48h, median denoted with vertical bar (orange). (**c**) Dot plots of fluorescence lifetime imaging redox ratio (FLIRR) colored as prior with each point representing single cell value, median denoted with vertical bar (orange). (**d**) Heatmap of OMI parameters corresponding to patient labels in (**a**) and (**b**) with color-coded z-score. Z-score defined using 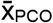 (average value of individual OMI parameter for an individual PCO culture), 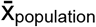 (average value of individual OMI parameter across the population), σ_population_ (standard deviation of an individual OMI parameter across the population). Distribution plots of FLIRR colored by cancer histology including (**e**) FLIRR, (**f**) NAD(P)H τ_m_, and (**g**) FAD τ_m_.

Across all 31 PCO lines, 30,877 single cells were analyzed using OMI (**Fig 3c**). The median FLIRR across populations (0.349) was not significantly exceeded by the median range within a given culture (0.400, p=0.16; **Fig 3c**). Across samples, the OMI parameters did not cluster by tumor grade, staging, or disease type by primary histology (**Fig 3d, Fig S5**). PCOs generated from alternative histologies including ovarian, lung, and gastrointestinal stromal tumors (GISTs), have OMI parameters that fall within the distributions of PCOs derived from CRC (**Fig 3e-g**).

### Subculturing of PCOs can detect rare driver alterations with potential clinical importance

A patient with familial adenomatous polyposis syndrome with a germline pathologic alteration in the *Adenomatous Polyposis Coli* gene (*APC*^Q789*^) had PCOs derived by multisite sampling of 4 polyps and 5 distinct regions of a single transverse colon cancer (**Fig 4a**). Pathologic alterations in *APC* were present in tissue from all polyp and cancer samples. Subclonal changes in normalized variant allele frequency (nVAF) were observed between patient tissue and the expanded PCOs, respectively (*APC*^*R499\**^ in polyp 1, nVAF 41% v. 89%; *APC*^*R554\**^ in polyp 4, nVAF 28% v. 49%; *APC*^*R1450\**^ in tumor 2, nVAF 0% v. 50%; and *BRAF*^*V600_K601delinsE*^ in Tumor 3, nVAF 0% v. 48%; **Fig 4b**). To clarify the contributions of individual PCOs in the diversity of subclonal populations, single PCOs were randomly selected for further individual clonal expansion (**Fig 4c**). Polyp 1 had persistent loss of *APC*^R499*^ across three independent PCO expansions. With expansion, polyp 2 harbored diversity in *APC* mutations, including the known germline *APC*^Q789*^, as well as *APC*^R499*\**^ which was not detected in the parent culture. Spikes from tumor 2 harbored *FBXW7*^R479Q^ (nVAF 49%) which was not detected by sequencing from either the parent culture of tumor 2, the expanded PCOs, or the primary tissue.

**Figure 4:**
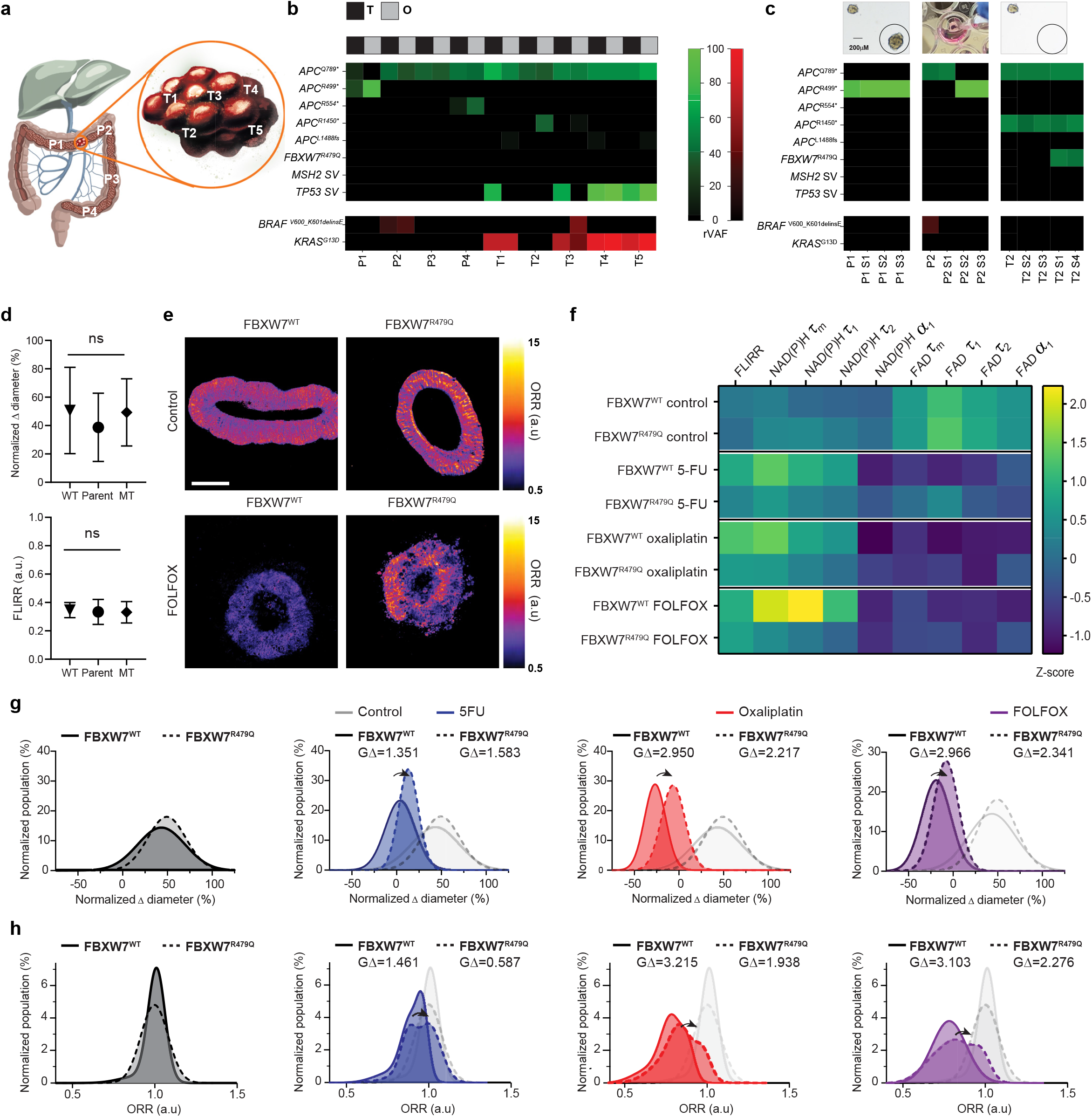
FOLFOX resistance identified in PCOs from a patient with familial adenomatous polyposis. (**a**) Sites of tissue sampling from patient with multiple site polyp (n = 4) and tumor sampling (n = 5). Heatmap of pathologic alterations of PCOs derived from individual polyps (P1-P4) and tumors (T1-T5) compared between primary tissue (black, T) and organoids (gray, O) plotted as relative variant allele frequency (rVAF). (**b**) Denoted are tumor suppressor genes (green) and oncogenes (red) plotted as a function of rVAR. (**c**) Heatmap of expanded PCO subclones selected by individual spikes using NGS profiling. (**d**) Comparison of normalized Δ diameter and FLIRR at 48h stratified by parent culture, *FBXW7*^WT^ (wild type, WT), and *FBXW7*^R479Q^ (mutant, MT) using two-sided student t-test (p > 0.05). (**e**) Representative PCOs metabolism assessed at 48h by ORR (NAD(P)H/FAD) for control (top panel) and FOLFOX stratified by *FBXW7* profile. Scale bar represents 100μm. (**f**) Heatmap of OMI parameters by *FBXW7* status stratified with respective Z-score as compared to parent culture. Z-score defined by 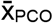 (average value of individual OMI parameter for individual PCO culture), 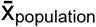 (average value of individual OMI parameter across the population), σ_population_ (standard deviation of an individual OMI parameter across the population). Gaussian distribution plots of normalized PCO diameter change assessed from 0 to 48h including control (gray), 5-FU (blue), oxaliplatin (red), and FOLFOX (purple). Molecular profile at *FBXW7* denoted wildtype (WT, solid line) and mutant *FBXW7*^R479Q^ (MT, dashed line). Response assessed using effect size (GΔ) relative to untreated control stratified by molecular profile at *FBXW7* for (**g**) normalized Δ diameter and (**h**) ORR at 48h.

Next, the treatment response to standard FOLFOX chemotherapy was compared between those organoids with and without the *FBXW7*^R479Q^ mutation to characterize the contributions of this isolated pathologic alteration. Relative to the parent population, the normalized change in diameter and FLIRR per replicate was consistent across subcultures (**Fig 4d**). There was not a detectable difference in the ORR between the *FBXW7*^WT^ and *FBXW7*^R479Q^ PCOs prior to treatment. After FOLFOX treatment the ORR was persistently higher for *FBXW7*^R479Q^ compared to *FBXW7*^WT^ PCOs, as the ORR of *FBXW7*^WT^ PCOs was reduced after FOLFOX treatment (**Fig 4e**). Further analysis of OMI parameters revealed enhanced response for NAD(P)H lifetime parameters τ_m_, τ_1_, and τ_2_ with FOLFOX treatment in *FBXW7*^R479Q^ cultures (**Fig 4f)**. Response as assessed by normalized change in diameter was similar for 5-FU alone with GΔ 1.4 (WT) v. 1.6 (MT). *FBXW7*^R479Q^ conferred reduced sensitivity for oxaliplatin with GΔ 3.0 (WT) v. 2.2 (MT) as well as FOLFOX with GΔ 3.0 (WT) v. 2.3 (MT) (**Fig 4g**). Consistent with changes in PCO diameter, metabolic response using ORR was reduced in *FBXW7*^R479Q^ PCOs for 5-FU (GΔ 1.5 v. 0.6), oxaliplatin (GΔ 3.2 v. 1.9), and FOLFOX (GΔ 3.1 v. 2.3) (**Fig 4h**).

### Organoid-level growth and single-cell OMI identify therapeutic sensitivity of CRC PCOs to EGFR inhibition

It is well-established in metastatic CRC that *K/NRAS* and *BRAF*^*V600*^ mutations lead to clinical therapeutic resistance to EGFR monoclonal antibodies, panitumumab and cetuximab. The activity of panitumumab was assessed in a panel of CRC PCOs with known mutation profiles (**Fig 5**). Population modeling after EGFR inhibition in MC4, possessing a *RAS*^A146V^ mutation, revealed two distinct populations both with persistent growth. In contrast, MR3 (*RAS*^WT^/*RAF*^WT^) PCOs converged to a single population that achieved growth arrest, with most organoids decreasing in size after EGFR inhibition (**Fig 5a-b**). Single-cell OMI did not demonstrate a change in the ORR following EGFR inhibition for MC4 (*RAS*^A146V^) as expected (**Fig 5c**). OMI of MR3 (*RAS*^WT^/*RAF*^WT^) revealed a shift in the ORR following panitumumab treatment (GΔ = 0.49; **Fig 5d**). The sensitivity of response was assayed across *RAS*^MT^ or *RAF*^MT^ PCOs (n = 4) and *RAS*^WT^/*RAF*^WT^ PCOs (n = 9). *RAS*^MT^ or *RAF*^MT^ PCOs had minimal changes in size between treatment (GΔ = 0.51, **Fig 5e**). In *RAS*^WT^/*RAF*^WT^ PCOs, measurements at 48h alone did not change significantly with treatment. The normalized change in diameter with individual organoid tracking across the population did identify a large effect size (GΔ = 1.3, **Fig 5f**), indicating the enhanced sensitivity of assessing treatment response on the individual organoid level over a treatment course. This was consistent with individual culture response across an expanded panel of PCOs, with significant differences in effect sizes observed when stratified by *RAS*/*RAF* status (p < 0.005; **Fig 5g**). No PCOs with *RAS*^*MT*^ or *RAF*^*MT*^ achieved an effect size ≥0.75, whereas the *RAS*^WT^/*RAF*^WT^ had variable effect size (GΔ 0.74–2.1). Clinically the patient’s cancer from which MR3 was derived was controlled with panitumumab at the site of the biopsy, which was used for PCO generation, however the patient had disease progression at distant sites with *KRAS* amplification (copy number = 89; **Fig 5h**).

**Figure 5:**
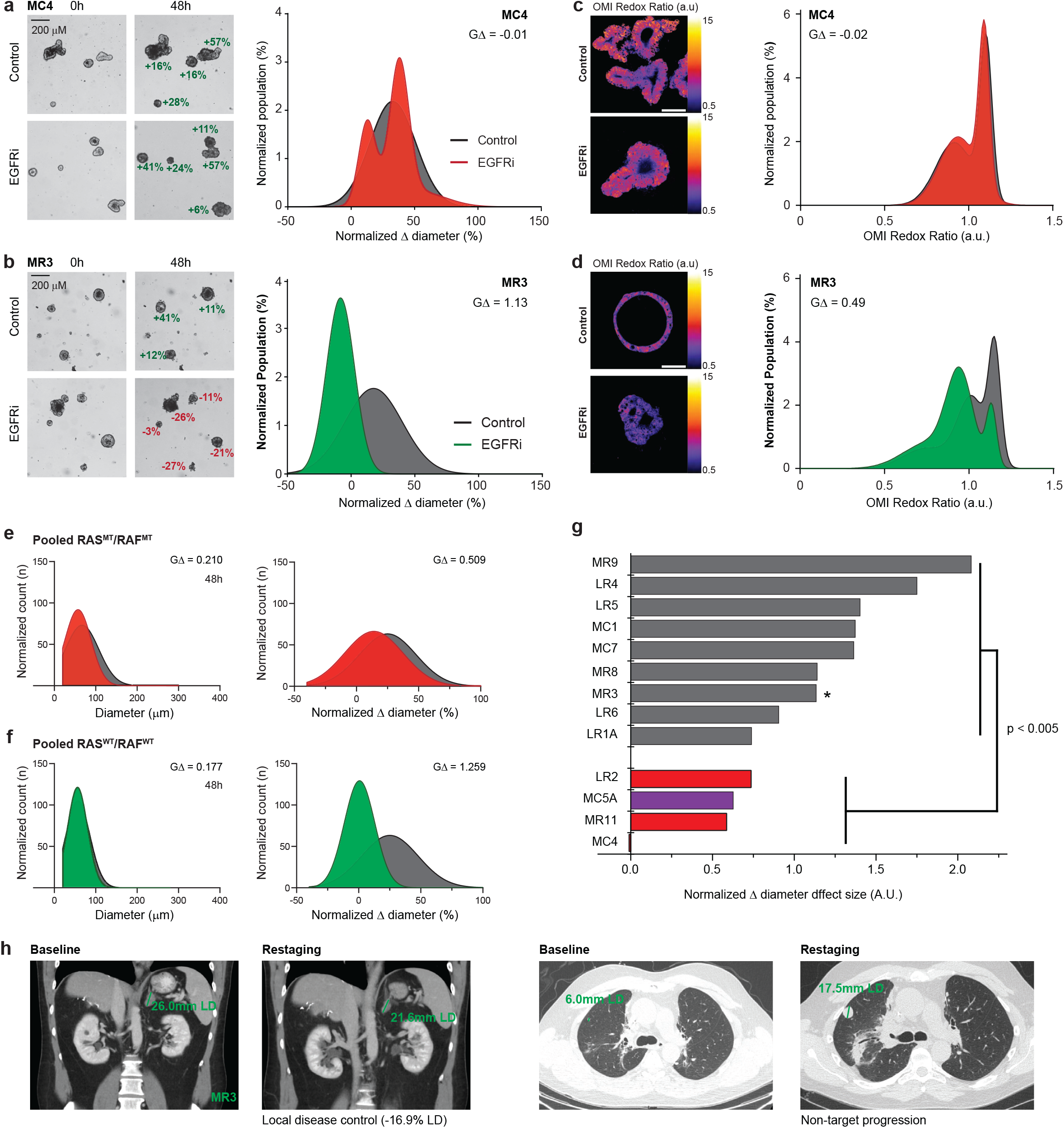
Assessment of PCO response to EGFR inhibition. (**a**) Representative brightfield images of therapeutic resistance of *KRAS*^A146V^ MC4 with persistent growth of control and panitumumab from 0h to 48h in contrast to (**b**) *RAS*^WT^ MR3 with growth arrest assessed by normalized Δ diameter at the organoid level. (**c**) Representative ORR for MC4 (*KRAS*^A146V^) and (**d**) MR3 (*KRAS*^WT^) with corresponding single cell distributions at 48 hours, scale bar represent 100μm. (**e**) Normalized ORR (NAD(P)H/FAD) shown in arbitrary units (a.u.). Pooled analysis of diameter for four independent lines predicted for resistance to EGFRi: *RAS*^MT^ (LR2, MR11, MC4) and *BRAF*^V600E^ (MC5A) at 48h (left panel) and change in diameter at 48h (right panel) by assessment of individual PCOs normalized to baseline diameter at 0h with corresponding effect size across distributions (GΔ). (**f**) Pooled analysis of diameter for nine independent *RAS*^WT^/*BRAF*^WT^ PCOs at 48h (left panel) and change in diameter at 48h (right panel) by assessment of individual PCOs normalized to baseline diameter at 0h with corresponding effect size across distributions (GΔ). (**g**) Line-specific sensitivity plotted by effect size (GΔ) including *RAS*^WT^/*BRAF*^WT^ (gray) compared against *RAS*^MT^ (red) and *BRAF*^V600E^ (violet) using student’s t-test for effect size of normalized Δ diameter. (**h**) MR3 PCO response of single agent EGFRi (panitumumab) (*denoted in **g**) represents a prospective clinical assessment tracked on CT scan at a 15-week follow-up showing local disease control at the biopsy site and a non-target progression in the right upper lung. Green lines indicate longest diameter (LD) of adrenal metastasis at baseline and restaging and measurements of non-target disease progression in right lung.

### PCOs response predicts clinical activity

Clinical response was prospectively correlated from PCOs developed from 13 patients with advanced cancers from the University of Wisconsin’s Precision Medicine Molecular Tumor Board and 4 controls with canonical mechanisms of resistance to EGFR inhibition (**Table S4**). The effect of single agent gemcitabine on MTB-3 (ovarian cancer) was negligible (**Fig 6a** and **b**; GΔ 0.40) consistent with a clinical outcome of progression in retroperitoneal adenopathy (**Fig 6c**). However, MTB-3 had intermediate sensitivity to single agent paclitaxel (GΔ = 1.1, **Fig 6b**). Metastatic CRC MC7 was collected at the time of clinical diagnosis with significant experimental response predicted to FOLFOX (GΔ = 2.2, **Fig 6d** and **e**) which matched the durable clinical response in the liver (**Fig 6f**).

**Figure 6.**
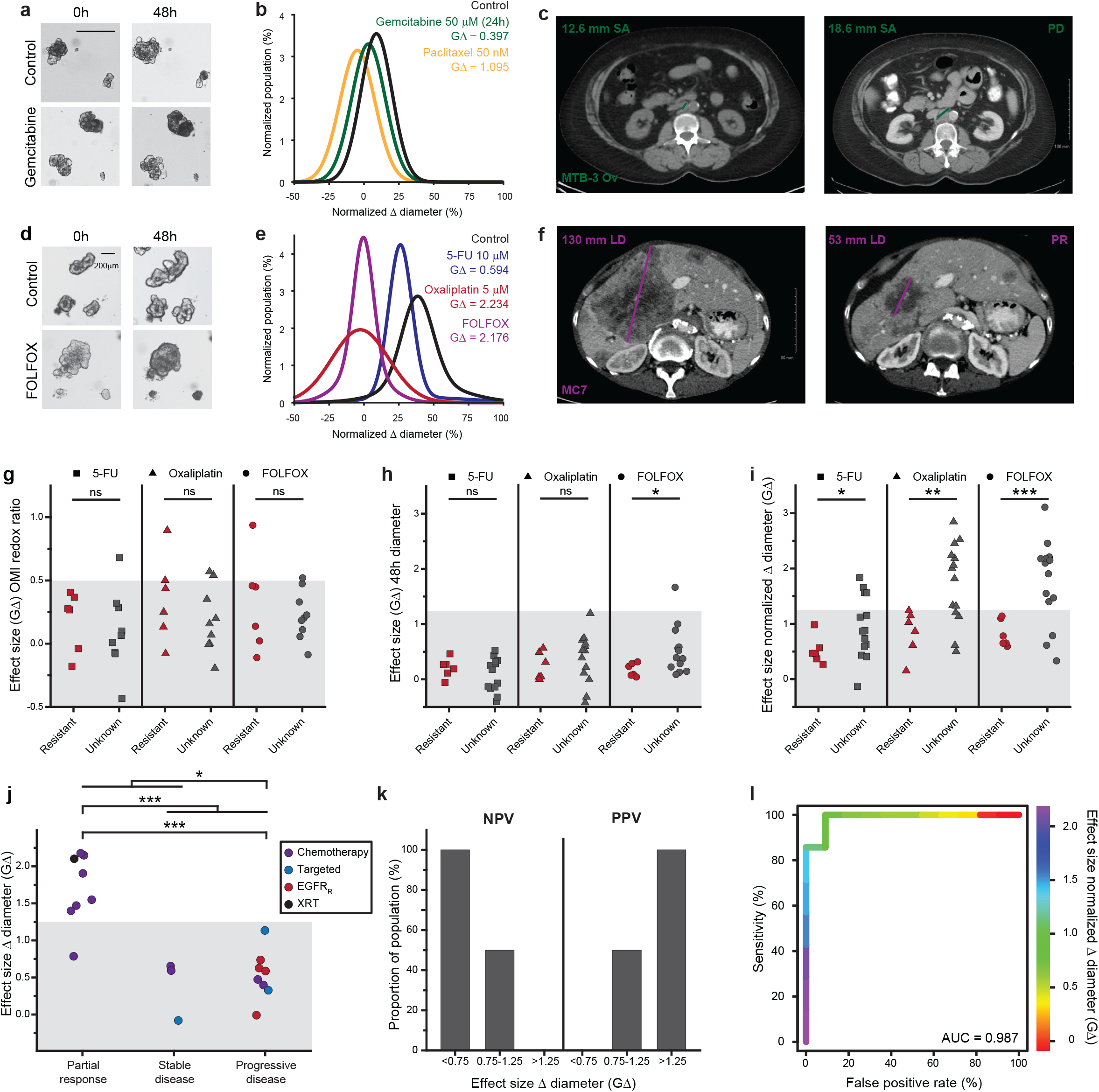
Validation of PCO response for clinical prediction. (**a**) Representative brightfield microscopy from MTB-3 ovarian (Ov) PCOs at baseline and 48h; scale bar represents 200μm for each panel. (**b**) Gaussian distributions for growth at 48 hours with respective effect sizes (GΔ) for MTB-3 Ov PCOs treated with gemcitabine 50μM (24 hours, green), paclitaxel 50nM (48 hours, gold) or control (black) as assessed at 48 hours. (**c**) Clinical response from the initial restaging CT scan of subject MTB-3 confirming the disease with enlarging retroperitoneal adenopathy after treatment with single agent gemcitabine on CT imaging. (**d**) Representative brightfield microscopy for MC7 PCOs treated with control (top panels), or FOLFOX (5-FU 10μm and oxaliplatin 5μm). (**e**) Gaussian distributions of MC7 for Δ diameter over 48h and respective effect sizes (GΔ) for 5-FU 10μm (blue), oxaliplatin 5μm (red), and FOLFOX (violet). (**f**) Restaging CT scan of MC7 shows partial response after FOLFOX. (**g-i**) Comparison of FOLFOX effect size (GΔ) between PCOs with disease progression after FOLFOX chemotherapy versus subjects without prior drug exposure assessed using two-sided student t-test with prior established sensitivity thresholds (shaded region). (**g**) Effect size of OMI redox ratio assessed at 48 hours was not significant (ns) between the clinically resistant and unknown cohorts. (**h**) Absolute diameter effect size assessed at 48 hours for single agent 5-FU (ns), oxaliplatin (ns) and FOLFOX (*p < 0.05) between clinically resistant and unknown cohorts. (**i**) Effect size of growth (percent Δ diameter) tracked from 0h to 48h for single agent 5-FU (*p < 0.05), oxaliplatin (**p < 0.005) and FOLFOX (***p < 0.0005) between clinically resistant and unknown cohorts. (**j**) Experimental sensitivity with clinical outcome or canonical mechanism of resistance labeled by treatment type including chemotherapy (purple), targeted therapy (blue), canonical EGFRi resistance (RAS^MT^ or RAF^MT^, red), and radiation (black) with reported significance by two-sided student t-test. (**k**) Bar plot of negative predictive value (NPV) and positive predictive value (PPV) for prospectively treated subjects. (**l**) Receiver operator curve (ROC) in response prediction plotted as false positive rate versus sensitivity with the colored line showing the continuum of effect size (GΔ) for change in diameter and corresponding area under the curve (AUC).

To identify thresholds for therapeutic resistance, we used PCOs with historic resistance to clinical treatments or otherwise unknown clinical response (**Fig 6g-i**). Five independent PCO lines were established from subjects with disease progression or recurrence after FOLFOX chemotherapy. After 48 hours of treatment, ORR lacked sensitivity in differentiating treatment response when compared to a panel of subjects without prior FOLFOX (**Fig 6g-i**). The effect size of 5-FU had modest association with baseline culture growth (R^2^ = 0.278) with improved correlation for oxaliplatin (R^2^ = 0.543) and FOLFOX (R^2^ = 0.560; **Fig S6**). In contrast to absolute diameter, the normalized change in diameter using individual organoid tracking improved sensitivity for identifying response to 5-FU (p < 0.05), oxaliplatin (p < 0.005) and FOLFOX (p < 0.0005; **Fig 6h**,**i**).

Previously published effect size thresholds were prospectively evaluated across PCOs comparing the PCO response to the RECIST v1.1 clinical response assessment for patients undergoing treatment with targeted therapy, cytotoxic chemotherapy, and/or radiotherapy^8^. PCO treatment response differentially predicted partial response versus progressive disease for most patients (**Fig 6j**). For those patients whose PCOs had a GΔ >1.25 in response to the treatment that they received clinically, a partial response (at least 30% reduction in cancer diameter) was observed in all patients (**Fig 6k**). Additionally, if the patient’s PCOs were limited to a GΔ <0.75, no partial responses to clinical treatment were observed (**Fig 6k**). 50% of the population for those patients whose PCOs had a GΔ between 0.75 and 1.25 achieved a partial response (**Fig 6k**). A receiver operator curve for partial response prediction achieved AUC of 0.987 as stratified by GΔ (**Fig 6l**).

### Longitudinal evaluation of PCO response to dose escalation of EGFR inhibitors in determining effiacy

As the growth and OMI evaluations do not require labels, dyes, or organoid destruction, these methods lend themselves to longitudinal PCO assessments. To improve the accessibility of these methods, epifluorescence wide-field microscopy was used to measure the ORR as panitumumab treatments were serially dose escalated based on culture growth (**Fig 7a**). As a representative example (MC7), the mean values of ORR were relatively stable during serial growth, with a decrease at 3 weeks and an increase at 6 weeks (**Fig 7b**). When visualized as normalized Gaussian distributions, divergent populations developed at week 3 (**Fig 7c**). The culture continued to achieve persistent growth at 100% physiologic C_max_ EGFRi which shifted to an increase in the ORR by week 6 of dose escalation (p<0.00005, **Fig 7b-c**).

**Figure 7.**
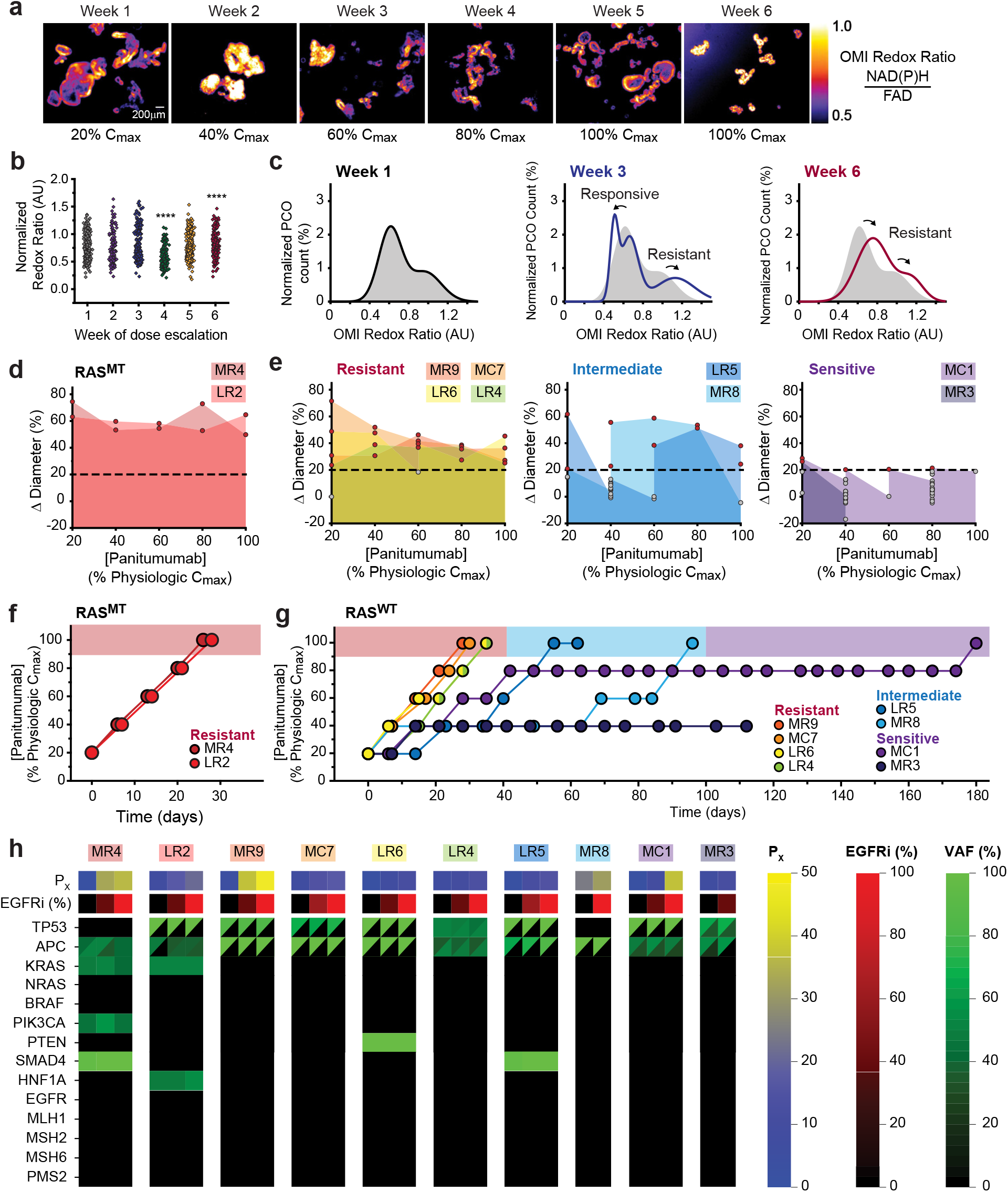
*Ex vivo* dose escalation of CRC PCOs. (**a**) Representative metabolic heterogeneity over the course of dose escalation of MC7 using epifluorescence microscopy for ORR (NAD(P)H/FAD) tracked weekly. (**b**) Sphere-level ORR for MC7 PCOs measured at 96h. All values normalized to week 1 mean signal intensity with points representing individual PCOs. ****p < 0.00005 as compared to redox ratio of week 1 dose escalation. (**c**) Normalized Gaussian distributions of individual PCO ORR for MC7 over a time course including week 1 (black line), week 3 (blue line), and week 6 (maroon line) plotted with the week 1 distribution (gray fill). (**d-e**) Median population growth rate tracked over the course of dose escalation as a percent of physiologic C_max_, labeled by growth rate with a threshold of growth (≥20%, red) versus insignificant growth (<20%, gray). Growth profiling over time course stratified by *RAS* mutation profile including (**d**) MT and (**e**) WT. (**f-g**)Time course of dose escalation stratified by *RAS* mutation profile including (**f**) MT and (**g**) WT. (**h**) Serial molecular profiling by cancer hotspot next generation sequencing of PCOs over the course of dose escalation with heat map labeling of absolute passage (P_x_, blue-yellow), physiologic C_max_ of EGFRi (black-red), and alteration variant allele frequency (VAF, black-green) with split cells (triangle) representing multiple gene-specific alterations.

The longitudinal tracking of resistance was objectively compared using the time to resistance (TTR), defined as the time to persistent growth at 100% physiologic C_max_ panitumumab, across ten CRC PCO lines. RAS^MT^ PCOs achieved serial thresholds for growth each week with *ex vivo* resistance determined within 30 days (**Fig 7d, f**). Across the RAS^WT^/RAF^WT^ PCOs a wide range in TTR was noted (**Fig 7e, g**). Primary resistance was identified in MR9, MC7, LR6 and LR4 with a TTR of <40 days. Additional lines had intermediate TTR (LR5, MR8) between 40-100 days, and prolonged sensitivity either achieving therapeutic resistance at >100 days (MC1) or arrested proliferation (MR3) (**Fig 7e, g)**.

### Transcriptional heterogeneity at ex vivo resistance to EGFR inhibition

To further evaluate the patient-specific mechanisms by which panitumumab conferred resistance, pairwise sequencing from baseline and resistant PCOs was performed. Repeat NGS comparing the baseline to the resistant cultures did not reveal the acquisition of pathologic alterations in *RAS, BRAF* or extracellular kinase alterations in *EGFR* which are commonly identified clinically (**Fig 7h**). Despite 5,587 individual transcripts with statistical differences in pairwise sets (p_adj_<0.05) only 90 (1.6%) were shared between resistant cultures (**Fig 8a**). Gene set variation analysis revealed heterogeneity in the absolute number of pathways significantly altered at resistance to EGFR inhibition (**Fig 8b**). The transcriptional regulation at resistance to EGFR inhibition harbored significant diversity in both individual transcripts and pathways (**Fig 8c-d**).

**Figure 8.**
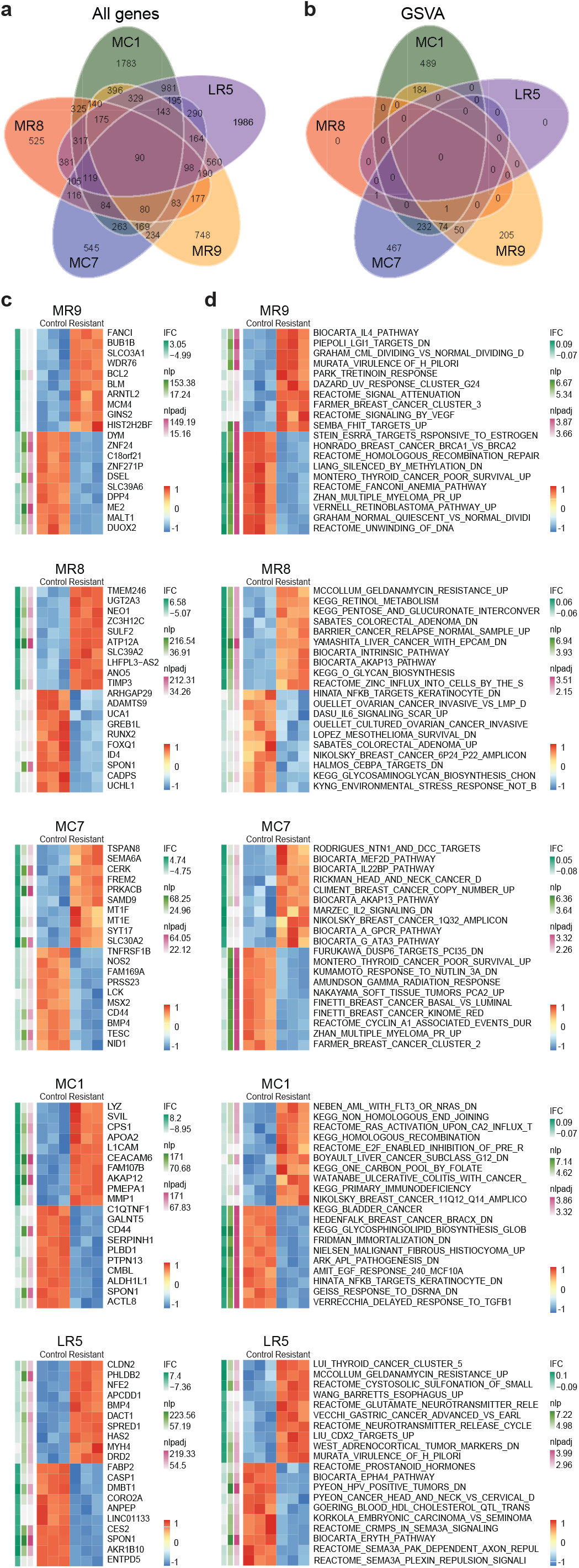
Transcriptional profiling at e*x vivo* resistant to EGFRi. (**a**) Venn diagram of RNASeq transcriptional profiles plotting significant differential expression (p_adj_ <0.05) from triplicate reads in pairwise sets between RAS^WT^/RAF^WT^ cultures and those expanded to EGFRi resistance. (**b**) Venn diagram of Gene Set Variation Analysis (GSVA) plotting the number of statistically significant altered pathways among RAS^WT^/RAF^WT^ cultures and those expanded to EGFRi resistance. Differential gene expression plotted as heatmaps for individual cultures at EGFRi resistance including (**c**) genes or (**d**) GSVA pathways plotted as top both up- and down-regulated expression.

## Discussion

There remains an urgent clinical need to develop personalized therapeutic strategies for patients with cancer. PCOs remain an exciting tool for understanding cancer biology and fostering drug development. This technology provides a potential mechanism for prediction of clinical response^8-10^. As more laboratories are expanding their use of PCOs for translational studies, detailed investigations are needed to identify the limitations of these models and how to best take advantage of the numerous strengths, including the representation of cancer cell heterogeneity for individual patients. Here we demonstrate PCO heterogeneity using a dedicated assessment of individual organoids tracked over time. Across a diverse set of CRC PCOs, baseline size, passage number, density, and location within the matrix did not confer differences in culture growth responsible for this heterogeneity. Novel multiphoton fluorescence lifetime imaging microscopy did not reveal a significant correlation between line specific metabolism and growth rate, indicating that growth and redox metabolism remain independent markers of cellular activity and serve complementary contributions in PCO assessment.

Taking the heterogeneity into consideration, we demonstrate the importance of therapeutic assessments using population modeling based on individual organoid tracking over time and single cell-level metabolic imaging. This method provides an alternative to ATP-based viability assays that are read using log-fold extensions of physiologic agents in defining therapeutic activity^24,47,9^. This is of particular importance as a sensitive technique to identify treatment response and sub-populations of PCOs with differential sensitivity. Additionally, PCOs hold great potential for the identification of important subclones with unique molecular alterations that could be important mechanisms of treatment resistance. The ability to subculture individual organoids to identify differences in the clonal structure of a given sample provides translational use such as defining contributions of select alterations in therapeutic sensitivity.

The use of PCOs in predicting clinical response should consider the benefit of modeling intratumoral heterogeneity. Using fixed, physiologic dosing, therapeutic response was assessed considering the potential for subclonal resistance across individual cultures. Alternative techniques, including traditional cell viability, yield non-viable cultures for which the characteristics of intermediate resistant cultures cannot be readily tracked. Population based modeling of response by change in individual PCO diameter was predictive of clinical response across a variety of clinical settings including the use of chemotherapy, targeted therapies, and radiation. Methods to track the dynamics of growth by normalizing baseline organoid culture parameters were integral for predicting clinical response in that assessment of organoid diameter alone failed to predict prior canonical sensitivities of resistance. These techniques can integrate with recent work to expand therapeutic response assessment using more accessible single photon technologies^78^.

Longitudinal time-course modeling in organoids provides many advantages. This technique was independently developed to overcome the lack of sensitivity to EGFR monoclonal antibodies reported in prior ATP-based viability assays^24^. TTR across these samples was of clinical relevance with serial resistance in all *RAS*^MT^ cultures, yet a range of sensitivities was observed in the *RAS*^WT^ cultures in a clinically relevant range of weeks to few months. Longitudinal evaluation of therapeutics using PCOs needs further clinical validation as a marker of duration of clinical response. Additionally, further studies are needed to determine if the mechanisms of resistance identified in PCOs also represent the mechanisms that occur clinically in the cancer from which the organoids are derived. It is of particular interest that the *RAS*^WT^/*RAF*^WT^ CRC PCOs were able to become resistant to anti-EGFR therapy in relatively short intervals with each of the investigated PCO lines becoming resistant for diverse reasons. This indicates the potential for PCOs in identifying patient specific resistance mechanisms, though it has yet to be seen if these studies can be performed in a clinically relevant time frame to make use of these mechanisms *ex vivo*. Of note, across these PCO lines investigated, we did not identify the acquisition of resistance mutations within our organoid cultures. Using circulating tumor DNA (ctDNA), it is quite common to identify the acquisition of mutations or other alterations in *RAS, RAF*, or *EGFR*, among others, though these are commonly at exceedingly low variant allele frequencies. The PCO data here calls to question the importance of these rare alterations in clinical resistance and could lead to future combined analyses with PCOs and ctDNA offering complementary data in identifying resistance mechanisms for individual patients.

This study details PCO response assessment to define the contributions of subclonal populations for clinical response prediction. The contributions of experimental parameters including organoid size, passage, and plating density did not predict differential growth across populations. Rather, it was line specific heterogeneity that could be readily selected using the expansion of individual PCOs, that persisted under therapeutic treatment, and that drove a prediction of clinical outcomes. Despite the promise of defining these unique populations, the complexities of transcriptional regulation are formidable with resistance including EGFR inhibition after dose escalation. This work provides a framework to characterize line specific adaptive mutability in response to targeted therapy^79^ and has been integrated into the prospective investigation of EGFRi for advanced CRC (NCT04587128).

## Acknowledgements

This project was supported by NIH grants R37CA226526 (DAD), T32 AG000213 (JDK), T32 CA009135 (KAJ), and P30 CA014520 (Core Grant, University of Wisconsin Carbone Cancer Center). The Skala laboratory is supported by grants from the NSF (CBET-1642287), Stand Up to Cancer (SU2C-AACR-IG-08-16, SU2CAACR-PS-18) and the NIH (R01 CA185747, R01 CA205101, R01 CA211082, U01 TR002383). JDK is supported by the Doris Duke Charitable Foundation’s Physician Scientist Fellowship and Conquer Cancer of ASCO’s Young Investigator Award. Additional support provided from Funk Out Cancer, the Cathy Wingert Colorectal Cancer Research Fund, and the ACI/Schwenn Family Professorship (DAD). David Latin Porter provided graphic support in preparation of this manuscript. The authors would like to thank members of the UWCCC GI DOT and the Translational Science BioCore and Experimental Pathology Shared Services which are supported by the UWCCC core grant from the NIH (P30 CA014520).

## Author Contributions

Conception and design: Jeremy D. Kratz, Peter F. Favreau, Cheri A. Pasch, Melissa C. Skala, Dustin A. Deming Development of methodology: Jeremy D. Kratz, Katherine A. Johnson, Peter F. Favreau, Cheri A. Pasch, Sean J. Mcilwain, Irene M. Ong, Melissa C. Skala, Dustin A. Deming

Acquisition of data: Jeremy D. Kratz, Shujah Rehman, Katherine A. Johnson, Amani A. Gillette, Aishwarya Sunil, Peter F. Favreau, Austin H. Yeung, Carley M. Sprackling

Analysis and interpretation of data (e.g., statistical analysis, biostatistics, computational analysis): Jeremy D. Kratz, Shujah Rehman, Katherine A. Johnson, Amani A. Gillette, Aishwarya Sunil, Peter F. Favreau, Cheri A. Pasch, Devon Miller, Lucas C. Zarling, Austin H. Yeung, Linda Clipson, Samantha J. Anderson, Alyssa K. Dezeeuw, Carley M. Sprackling, Sean J. Mcilwain, Irene M. Ong, Melissa C. Skala, Dustin A. Deming

Writing, review, and/or revision of the manuscript: Jeremy D. Kratz, Shujah Rehman, Katherine A. Johnson, Amani A. Gillette, Aishwarya Sunil, Peter F. Favreau, Cheri A. Pasch, Devon Miller, Lucas C. Zarling, Austin H. Yeung, Linda Clipson, Samantha J. Anderson, Alyssa K. DeZeeuw, Carley M. Sprackling, Kayla K. Lemmon, Daniel E. Abbott, Mark E. Burkard, Michael F. Bassetti, Jens C. Eickhoff, Eugene F. Foley, Charles P. Heise, Randall J. Kimple, Elise H. Lawson, Noelle K. LoConte, Sam J. Lubner, Daniel L. Mulkerin, Kristina A. Matkowskyj, Cristina B. Sanger, Nataliya V. Uboha, Sean J. Mcilwain, Irene M. Ong, Evie H. Carchman, Melissa C. Skala, Dustin A. Deming

Administrative, technical, or material support (i.e., reporting or organizing data, constructing databases): Jeremy D. Kratz, Shujah Rehman, Katherine A. Johnson, Amani A. Gillette, Linda Clipson, Kristina A. Matkowskyj, Sean J. Mcilwain

Subject Recruitment: Jeremy D. Kratz, Kayla K. Lemmon, Daniel E. Abbott, Mark E. Burkard, Michael F. Bassetti, Jens C. Eickhoff, Eugene F. Foley, Charles P. Heise, Randall J. Kimple, Elise H. Lawson, Noelle K. LoConte, Sam J. Lubner, Daniel L. Mulkerin, Kristina A. Matkowskyj, Cristina B. Sanger, Nataliya V. Uboha, Evie H. Carchman, Dustin A. Deming

Study supervision: Melissa C. Skala, Dustin A. Deming

