## Supplemental information for "Integrating Subclonal Response Heterogeneity to Define Cancer Organoid Therapeutic Sensitivity"

**Extended Data and Supplemental Figures**

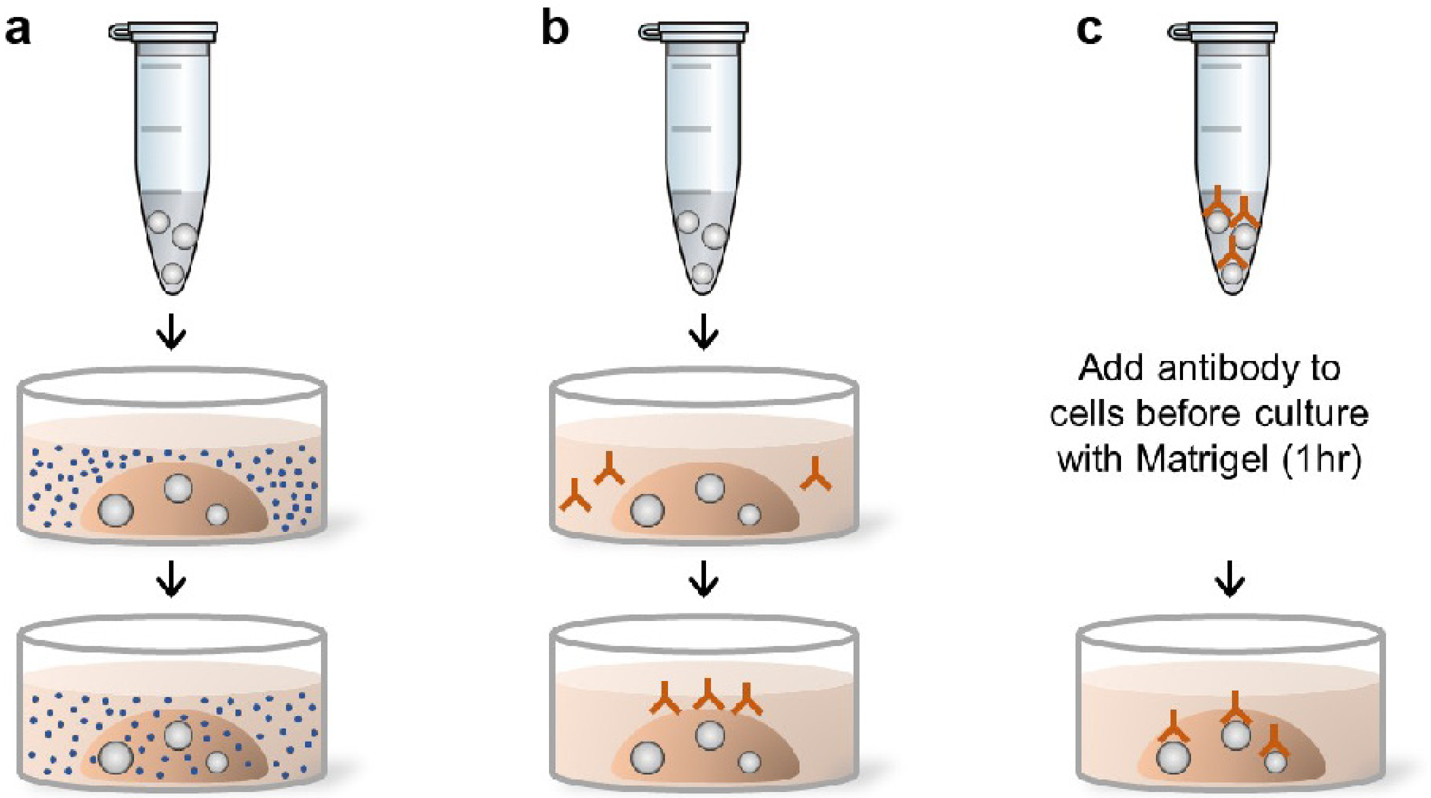


**Figure S1:** (**a**) Model for therapeutic delivery used in EGFR inhibitor assay design. Free diffusion assumed with small molecular delivery. (**b**) Efficient monoclonal antibody delivery through a 3D matrix was uncertain. (**c**) Assay design with 1hr incubation of cell suspension with panitumumab prior to plating.

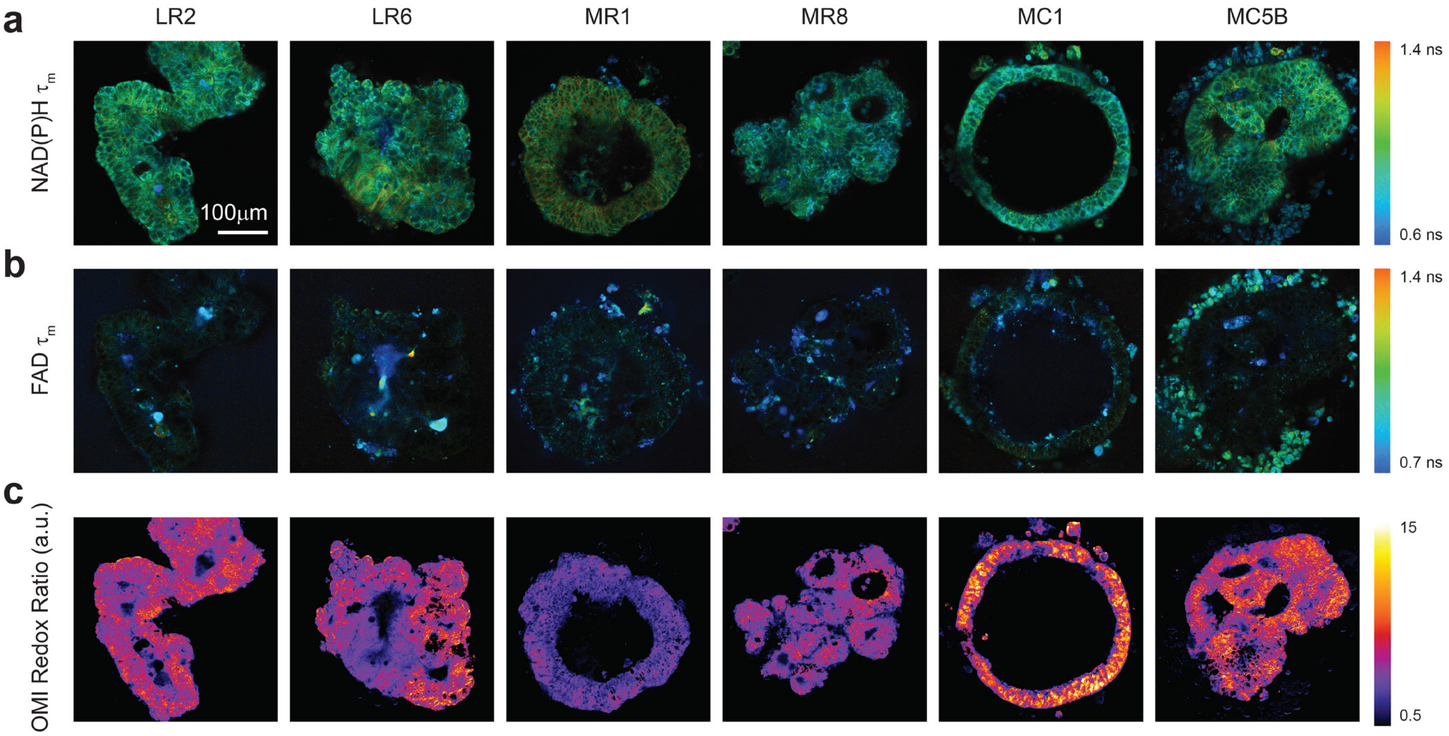


**Figure S2:** Representative examples of intra-organoid heterogeneity using OMI microscopy. (**a**) NADH(P)H and (**b**) FAD mean lifetime in nanoseconds (ns) of untreated organoids with corresponding (**c**) ORR (ratio of NAD(P)H/FAD) for single PCOs with normalization of arbitrary units (a.u.).

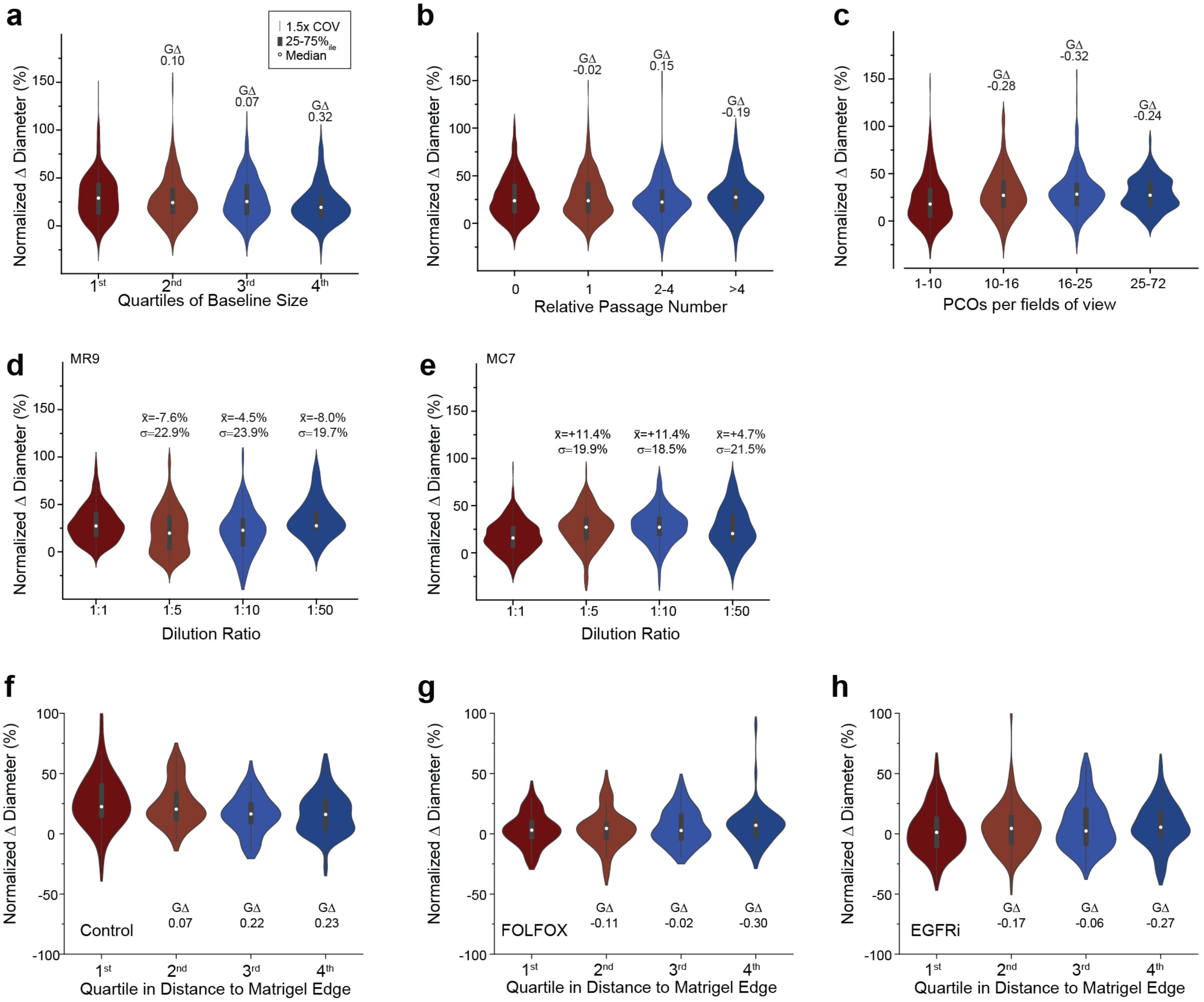


**Figure S3:** Pooled analyses of experimental contributions to growth profiling for Δ diameter plotted by interquartile populations of (**a**) baseline size, (**b**) relative passage number, and (**c**) number of PCOs per field of view. Contributions of plating density (dilution ratio) for control growth including mean growth ($\bar{x}$) and population standard deviation ($\sigma$) for (**d**) MR9 and (**e**) MC7. Pooled analysis of growth by interquartile populations of location relative to matrix edge for three independent lines for (**f**) control, (**g**) FOLFOX, and (**h**) EGFRi with corresponding effect size (GΔ). Plotted are 1.5x coefficient of variance (gray line), interquartile range (25-75%, gray box), and median (white dot) for each population).

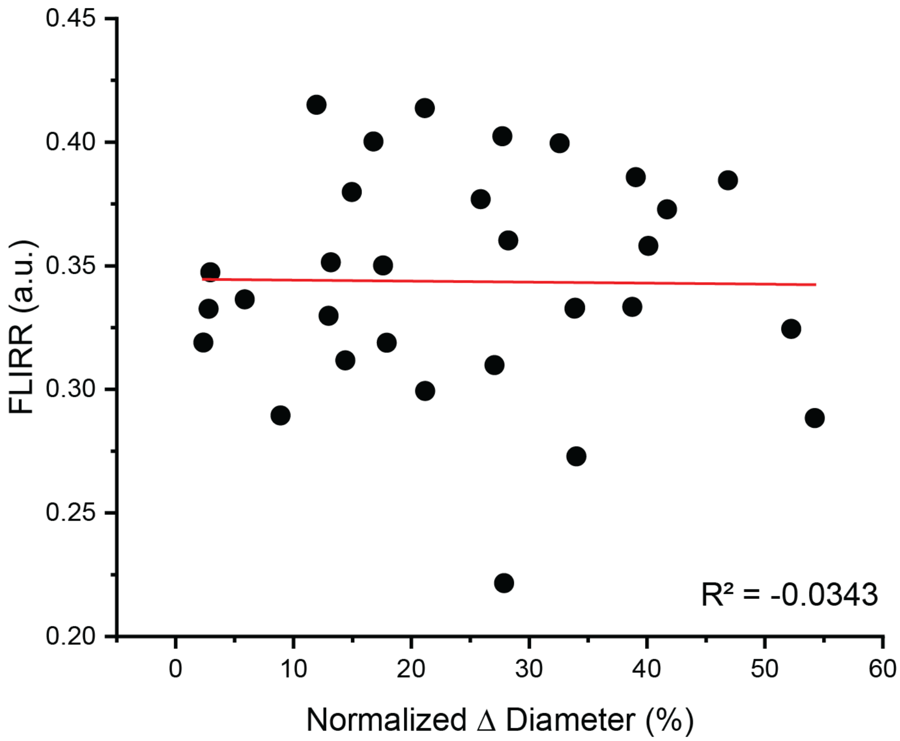


**Figure S4:** Correlation between median Δ diameter (0 to 48h) and median fluorescence lifetime redox ratio (FLIRR) at 48h between PCO lines (n=31) with corresponding adjusted correlation coefficient ($R^2$).

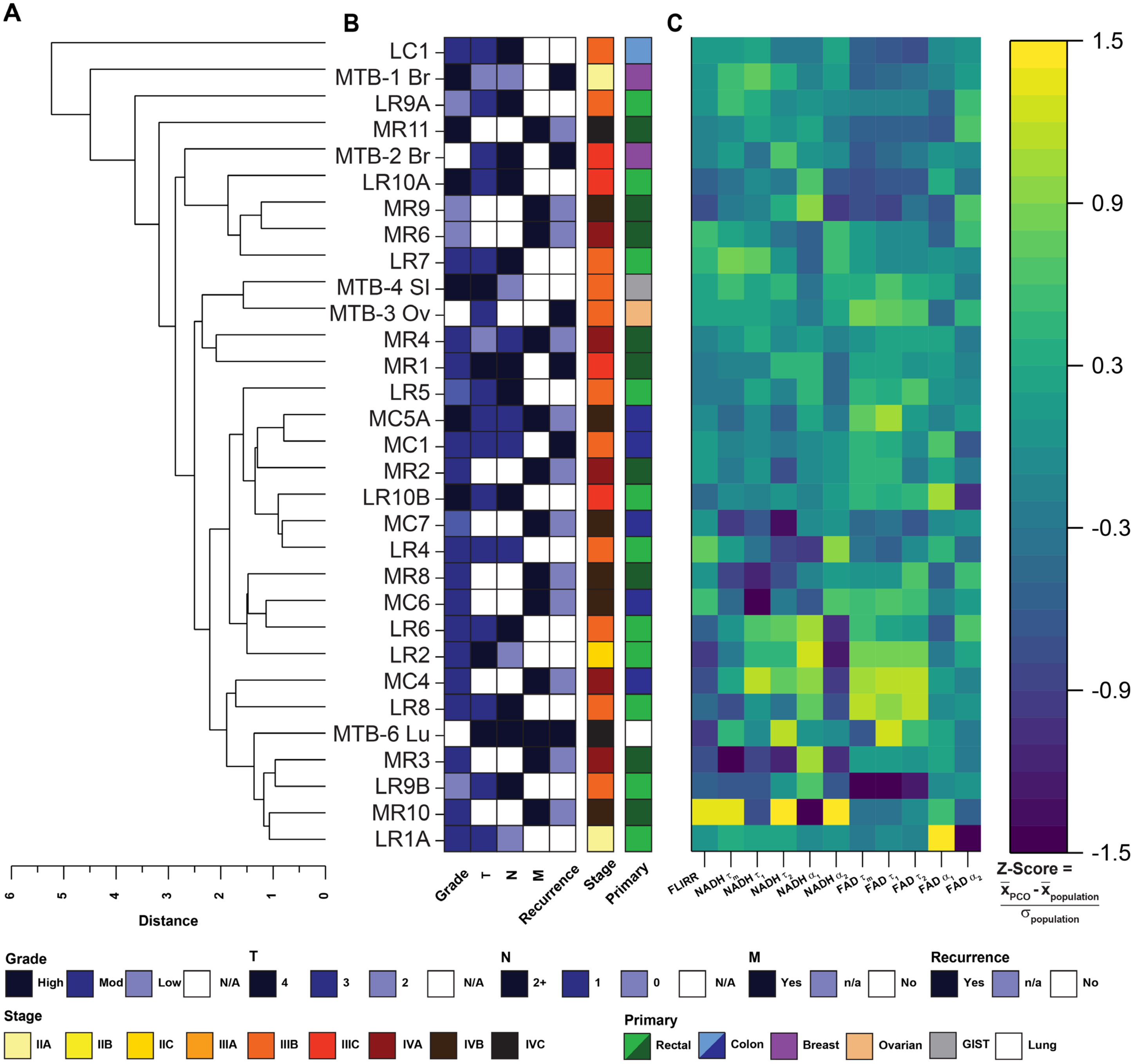


**Figure S5:** Hierarchical cluster analysis of OMI parameters. Dendrogram from cluster analysis with corresponding euclidean distance (**a**). Clinical characteristics for cultures colored by the key below, including pathologic review of tumor grade and clinical parameters of stage from tumor (T), node (N) and metastases (M) combined to complete diagnostic staging (Stage). Current disease status noted (recurrence in black). Primary cancer denoted by colored key of primary tumor type (Primary) for localized rectal (light green), metastatic rectal (dark green), localized colon (light blue), metastatic colon (dark blue), breast (purple), ovarian (beige), gastrointestinal stromal tumor (GIST, gray), and lung (white) (**b**). Heatmap of OMI parameters corresponding to patient labels in (**b**) with color-coded z-score (**c**). Z-score defined using $\bar{x}_{PCO}$ (average value of an individual OMI parameter for an individual PCO culture), $\bar{x}_{population}$ (average value of an individual OMI parameter across the population), $\sigma_{population}$ (standard deviation of an individual OMI parameter across the population).

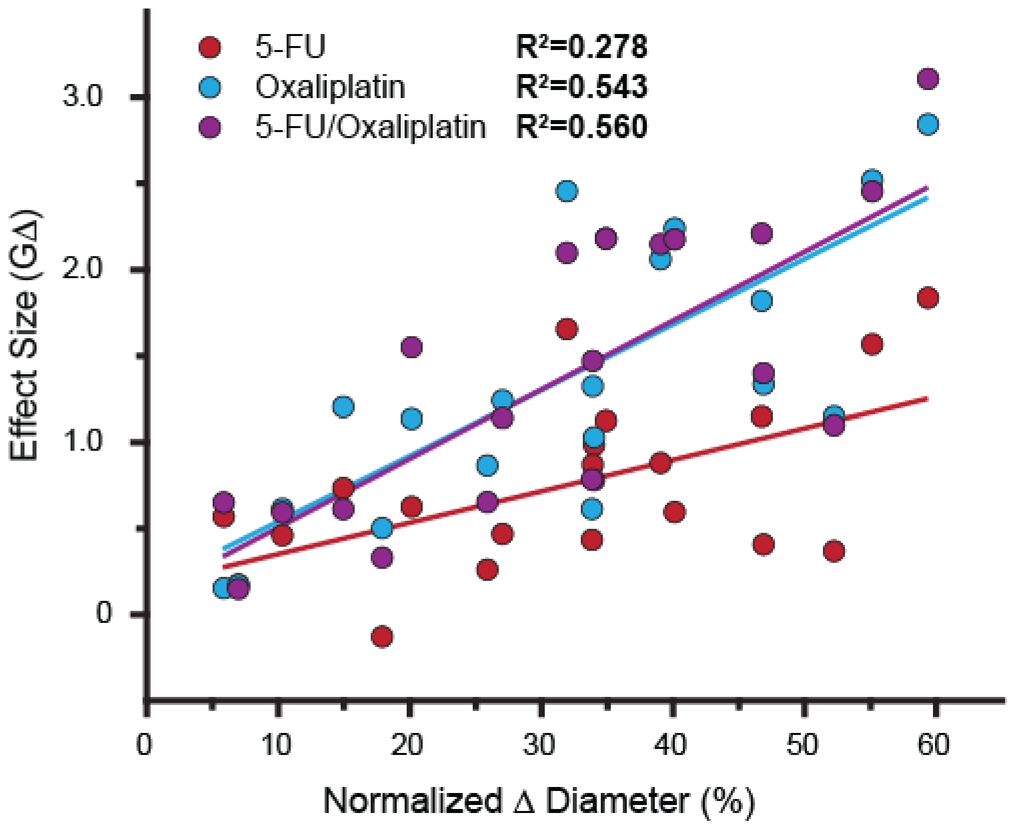


**Figure S6:** Correlation between mean normalized change (Δ) in diameter for individual PCOs plotted against the effect size of therapeutic sensitivity of the treatment groups including single agent 5-FU (red), oxaliplatin (blue), and combination FOLFOX (violet). Correlation co-efficient calculated using adjusted $R^2$.

**Table S1:** Summary of culture conditions used in maintenance and feeding of PCOs.

| Name | Component | Supplier & Catalog | Final Concentration | Application |
|---|---|---|---|---|
| **DMEM stock** | Advanced DMEM/F12 | Invitrogen 12634-100 | 1x | Digestion |
| | Fetal Bovine Serum | Sigma F8067-500ml | 10% | |
| | Pen/Streptomycin | Invitrogen 15140-122 | 100 IU/mL | |
| **ADF stock** | Advanced DMEM/F12 | Invitrogen 12634-100 | 1x | Washing, passaging |
| | Glutamax, 100x | Invitrogen 35050-061 | 1x | |
| | HEPES | Fisher SH30237.01 | 10mM | |
| | Pen/Streptomycin | Invitrogen 15140-122 | 50 IU/mL | |
| **ADF feed** | ADF stock | (as above) | 100% v/v | Breast, Ovarian, Lung, GIST |
| | EGF, recombinant mouse | Invitrogen 53003-018 | 50ng/mL | |
| **WNT feed** | ADF stock | (as above) | 50% v/v | Colon, Rectal |
| | WNT3a conditioned media | (in house) | 50% v/v | |
| | EGF, recombinant mouse | Invitrogen 53003-018 | 50ng/mL | |

**Table S2:** Summary of agents used in therapeutic studies.

| Agent | | Concentration | Duration | Vendor |
|---|---|---|---|---|
| FOLFOX | A) 5-FU | 10 μM | 48h | Fresenius |
| | B) Oxaliplatin | 5 μM | | Hospira |
| FOLFIRI | A) 5-FU | 10 μM | 48h | Fresenius |
| | B) SN-38 | 1.5 nM | | Sigma |
| Gemcitabine | | 50 μM | 24h | Hospira |
| Paclitaxel | | 50 nM | 48h | Athenex |
| Olaparib | | 200 nM | 48h | LC Laboratories |
| Osimertinib | | 30 nM | 48h | LC Laboratories |
| Panitumumab | | 1.6 μM | 48h | Amgen |

**Table S3:** Summary of PCOs in a therapeutic investigation organized by clinical stage including localized rectal (LR), metastatic rectal (MR), localized colon (LC) and metastatic colon (MC) as well as tissue acquired from the molecular tumor board including breast (Br), Ovarian (Ov), Small Intestine (SI), and Lung (Lu).

| Name | Histology | Primary Tumor | Site of Tissue | Tissue Sampling | Grade | T | N | M | Stage | Recurrence |
|---|---|---|---|---|---|---|---|---|---|---|
| LR1A | Adenocarcinoma | Rectum | Rectum | Surgical Resection | Moderate | pT3 | pN0 | M0 | IIA | No |
| LR1B | Adenocarcinoma | Rectum | Rectum | Surgical Resection | Moderate | pT3 | pN0 | M0 | IIA | No |
| LR2 | Adenocarcinoma | Rectum | Rectum | Surgical Resection | Moderate | pT4b | pN0 | M0 | IIC | No |
| LR3 | Adenocarcinoma | Rectum | Rectum | Surgical Resection | Moderate | T4 | N0 | M0 | IIC | No |
| LR4 | Adenocarcinoma | Rectum | Rectum | Endoscopic Biopsy | Moderate | T3b | N1b | M0 | IIIB | No |
| LR5 | Adenocarcinoma | Rectum | Rectum | Endoscopic Biopsy | Low-Moderate | T3 | N2a | M0 | IIIB | No |
| LR6 | Adenocarcinoma | Rectum | Rectum | Endoscopic Biopsy | Moderate | T3 | N2a | M0 | IIIB | No |
| LR7 | Adenocarcinoma | Rectum | Rectum | Endoscopic Biopsy | Moderate | T3c | N2a | M0 | IIIB | No |
| LR8 | Adenocarcinoma | Rectum | Rectum | Endoscopic Biopsy | Moderate | T3c | N2a | M0 | IIIB | No |
| LR9A | Adenocarcinoma | Rectum | Rectum | Endoscopic Biopsy | Low | T3a | N2a | M0 | IIIB | No |
| LR9B | | | | | | | | | | No |
| LR10A | Adenocarcinoma | Rectum | Rectum | Endoscopic Biopsy | High | T3c | N2 | M0 | IIIC | No |
| LR10B | | | | | | | | | | No |
| LR11 | Adenocarcinoma | Rectum | Rectum | Endoscopic Biopsy | High | T3 | N2b | M0 | IIIC | No |
| MR1 | Mucinous Adenocarcinoma | Rectum | Peritoneum | Surgical Resection | Moderate | pT4 | pN2a | M0 | IIIC | Yes |
| MR2 | Adenocarcinoma | Rectum | Liver | Surgical Resection | Moderate | n/a | n/a | M1a | IVA | N/A |
| MR3 | Adenocarcinoma | Rectum | Lung | IR Biopsy | Moderate | n/a | n/a | M1a | IVA | N/A |
| MR4 | Adenocarcinoma | Rectum | Liver | Surgical Resection | Moderate | pT2 | pN1b | M1a | IVA | N/A |
| MR5 | Adenocarcinoma | Rectum | Liver | Surgical Resection | Moderate | n/a | n/a | M1a | IVA | N/A |
| MR6 | Adenocarcinoma | Rectum | Liver | Surgical Resection | Low | n/a | n/a | M1a | IVA | N/A |
| MR7 | Adenocarcinoma | Rectum | Retroperitoneal Node | IR Biopsy | n/a | n/a | n/a | M1a | IVA | N/A |
| MR8 | Adenocarcinoma | Rectum | Liver | Surgical Resection | Moderate | n/a | n/a | M1b | IVB | N/A |
| MR9 | Adenocarcinoma | Rectum | Liver | IR Biopsy | Low | n/a | n/a | M1b | IVB | N/A |
| MR10 | Adenocarcinoma | Rectum | Rectum | Endoscopic Biopsy | Moderate | n/a | n/a | M1b | IVB | N/A |
| MR11 | Mucinous Adenocarcinoma | Rectum | Omentum | Surgical Resection | High | n/a | n/a | M1c | IVC | N/A |
| LC1 | Adenocarcinoma | Colon | Colon | Surgical Resection | Moderate | pT3 | pN2a | M0 | IIIB | No |
| LC2 | Adenocarcinoma | Colon | Colon | Surgical Resection | Moderate | pT4a | pN2b | M0 | IIIC | No |
| MC1 | Adenocarcinoma | Colon | Liver | Surgical Resection | Moderate | pT3 | pN1b | M0 | IIIB | Yes |
| MC2 | Mucinous Adenocarcinoma | Colon | Soft Tissue Mass | Surgical Resection | High | pT4b | pN1b | M0 | IIIB | Yes |
| MC3 | Adenocarcinoma | Colon | Liver | Surgical Resection | Low | pT3 | pN1a | pM1a | IVA | N/A |
| MC4 | Adenocarcinoma | Colon | Colon | Surgical Resection | Moderate | n/a | n/a | M1a | IVA | N/A |
| MC5A | Adenocarcinoma | Colon | Colon | Endoscopic Biopsy | Low-Moderate | pT3 | pN1a | M1b | IVB | N/A |
| MC5B | Adenocarcinoma | Colon | Colon | Surgical Resection | Moderate | pT3 | pN1 | M1a | IVB | N/A |
| MC6 | Adenocarcinoma | Colon | Colon | Surgical Resection | High | n/a | n/a | M1b | IVB | N/A |
| MC7 | Adenocarcinoma | Colon | Colon | Endoscopic Biopsy | Moderate | n/a | n/a | M1b | IVB | N/A |
| **Precision Medicine Molecular Tumor Board** | | | | | | | | | | |
| MTB-1 Br | ER-/PR-/Her2- Adenocarcinoma | Breast | Pericardial Effusion | Pericardiocentesis | High | T2 | N0 | M0 | IIA | Yes |
| MTB-2 Br | Infiltrating ductal carcinoma | Breast | Peritoneal Fluid | Paracentesis | NR | pT3 | pN3a | M0 | IIIC | Yes |
| MTB-3 Ov | Serous Carcinoma | Ovarian | Ovary | IR Biopsy | NR | pT3b | pNX | M0 | IIIB | Yes |
| MTB-4 SI | Spindle cell GIST | Small Intestine | Pancreas | Surgical Resection | High | pT4 | pN0 | M0 | IIIB | No |
| MTB-5 SI | Spindle cell GIST | Small Intestine | Small Intestine | Surgical Resection | Low | n/a | n/a | M1 | IV | N/A |
| MTB-6 Lu | Adenocarcinoma | Lung | Pleural Effusion | Thoracentesis | NR | T4 | N2 | M1c | IVC | Yes |

**Table S4:** Summary of prospectively tracked PCOs with corresponding response as assessed by effect size using Glass's Delta for change in diameter between control and respective therapy. All clinical responses were assessed per RECIST v.1.1 including partial response (PR), stable disease (SD), progressive disease (PD) or canonical mechanisms of EGFR inhibition with activating alterations in RAS/RAF.

| **Line** | **Therapy** | **Effect Size (G$\Delta$)** | **Clinical Response** | **RAS$^{MT}$/RAF$^{MT}$ EGFRi Reistance** |
|---|---|---|---|---|
| LR2 | Panitumumab | 0.736 | n/a | Yes |
| LR4 | FOLFOX | 1.549 | PR | n/a |
| LR5 | FOLFOX | 0.653 | SD | n/a |
| LR7 | FOLFOX | 2.148 | PR | n/a |
| LR9A | FOLFOX | 1.399 | PR | n/a |
| LR9B | FOLFOX | 1.470 | PR | n/a |
| LR10A | XRT | 2.101 | PR | n/a |
| MR10 | FOLFOX | 0.785 | PR | n/a |
| MR11 | Panitumumab | 0.586 | n/a | Yes |
| MR3 | Panitumumab | 1.134 | PD | n/a |
| MC4 | Panitumumab | -0.010 | n/a | Yes |
| MC5A | Panitumumab | 0.626 | n/a | Yes |
| MC5B | FOLFOX | 0.592 | SD | n/a |
| MC6 | FOLFIRI | 0.473 | PD | n/a |
| MC7 | FOLFOX | 2.176 | PR | n/a |
| MTB-1 Br | Olaparib | -0.081 | SD | n/a |
| MTB-3 Ov | Gemcitabine | 0.397 | PD | n/a |
| MTB-6 Lu | Osimertinib | 0.327 | PD | n/a |
